# A nitrogenase-like enzyme is involved in the novel anaerobic assimilation pathway of a sulfonate, isethionate, in the photosynthetic bacterium *Rhodobacter capsulatus*

**DOI:** 10.1101/2024.05.19.594900

**Authors:** Yoshiki Morimoto, Kazuma Uesaka, Yuichi Fujita, Haruki Yamamoto

**Affiliations:** Graduate School of Bioagricultural Sciences, Nagoya University, Japan

## Abstract

Prokaryotes contribute to the global sulfur cycle by using diverse sulfur compounds as sulfur sources or electron acceptors. Here we report that a nitrogenase-like enzyme (NFL) and a radical SAM enzyme (RSE) are involved in the novel anaerobic assimilation pathway of a sulfonate, isethionate, in the photosynthetic bacterium *Rhodobacter capsulatus*. The *nflHDK* genes for NFL are localized at a locus containing genes for known sulfonate metabolism in the genome. A gene *nflB* encoding an RSE is present just upstream of *nflH*, forming a small gene cluster *nflBHDK*. Mutants lacking any *nflBHDK* genes lost the ability to grow with isethionate as the sole sulfur source under anaerobic photosynthetic conditions, indicating that all four NflBHDK proteins are essential for the isethionate assimilation pathway. Heterologous expression of the *islAB* genes encoding a known isethionate lyase that degrades isethionate to sulfite and acetaldehyde restored the isethionate-dependent growth of a mutant lacking *nflDK*, indicating that the enzyme encoding *nflBHDK* is involved in an isethionate assimilation reaction to release sulfite. Furthermore, heterologous expression of *nflBHDK* and *ssuCAB* encoding an isethionate transporter in the closely related species *R. sphaeroides*, which does not have *nflBHDK* and cannot grow with isethionate as the sole sulfur source, conferred isethionate-dependent growth ability to this species. We propose to rename *nflBHDK* as *isrBHDK* (<u>is</u>ethionate reductase). The *isrBHDK* genes are widely distributed among various prokaryote phyla. Discovery of the isethionate assimilation pathway by IsrBHDK provides a missing piece for the anaerobic sulfur cycle and for understanding the evolution of ancient sulfur metabolism.

**Importance:** Nitrogenase is an important enzyme found in prokaryotes that reduces atmospheric nitrogen to ammonia and plays a fundamental role in the global nitrogen cycle. It has been noted that nitrogenase-like enzymes (NFLs), which share an evolutionary origin with nitrogenase, have evolved to catalyze diverse reactions such as chlorophyll biosynthesis (photosynthesis), coenzyme F_430_ biosynthesis (methanogenesis), and methionine biosynthesis. In this study, we discovered that an NFL with unknown function in the photosynthetic bacterium *Rhodobacter capsulatus* is a novel isethionate reductase (Isr), which catalyzes the assimilatory degradation of isethionate, a sulfonate, releasing sulfite used as the sulfur source under anaerobic conditions. Isr is widely distributed among various bacterial phyla, including intestinal bacteria, and is presumed to play an important role in sulfur metabolism in anaerobic environments such as animal guts and microbial mats. This finding provides a clue for understanding ancient metabolism that evolved under anaerobic environments at the dawn of life.

## Introduction

Nitrogenase is a metalloenzyme responsible for biological nitrogen fixation, catalyzing the reduction of atmospheric nitrogen molecules (N_2_) to ammonia (NH_3_), which provides a nitrogen source to support most organisms and plays a critical role in the global nitrogen cycle [1]. Nitrogenase comprises three subunits (NifH, NifD, and NifK) [2, 3]. NifH functions as the reductase component as a homodimer (Fe protein), and NifD and NifK form a heterotetramer (MoFe protein) to function as the catalytic component. Nitrogenase-like enzymes (*nif*-like enzymes; NFLs) are a group of metalloenzymes that share high structural similarity with nitrogenase and catalyze a variety of reductions different from the conversion of N_2_ to NH_3_ [4–6]. NFLs include NifEN, which is involved in the biosynthesis of the metal center (FeMo-co) of nitrogenase [3, 7, 8]; dark-operative protochlorophyllide oxidoreductase (DPOR; BchLNB/ChlLNB) [9–11] and a chlorophyllide *a* oxidoreductase (COR; BchXYZ) [12], which are involved in (bacterio)chlorophyll biosynthesis; Ni^2+^-sirohydrochlorin *a*,*c*-diamide reductive cyclase (CfbCD), which is involved in the biosynthesis of the cofactor F_430_ essential for the anaerobic oxidation of methane in methanotrophs and the methane formation in methanogens [13, 14]; and methylthioalkane reductase (MarHDK), which is involved in methionine biosynthesis [15]. The NFLs consist of three subunits homologous to NifH, NifD, and NifK (only Cfb lacks a NifK homolog) of which the NifH homologs (NifH, BchL/ChlL, BchX, CfbC, MarH) function as the reductase components, and the NifD and NifK homologs (NifEN, BchNB/ChlNB, CfbD, MarDK) form a heterotetramer (only CfbD is a homodimer) as the catalytic components to catalyze the reduction of their individual substrates using electrons from the NifH homologs. The reductase component, the NifH homolog, carries a [4Fe-4S] cluster that serves as the redox center for electron transfer to the cognate catalytic component [3, 14, 16–19]. The catalytic component carries at least a pair of [4Fe-4S] clusters, and the electrons from the reductase component are used to reduce the substrate via the [4Fe-4S] clusters [3, 11, 14, 20, 21]. Thus, nitrogenase and NFLs catalyze various reductions through inter- and intramolecular electron transfers based on a common architecture with their metal clusters as the redox centers. These metal clusters, particularly the [4Fe-4S] cluster of the reductase component, are rapidly and irreversibly destroyed by oxygen (O_2_) [3, 19, 22]. Therefore, an anaerobic environment is required for the action of NFLs, similar to nitrogenase.

NFL genes are distributed among various prokaryotes. Raymond et al. (2004) classified NFLs into five groups (I–V) on the basis of molecular phylogenetic analysis [4]. Three types (Mo, V, and Fe) of nitrogenases were classified into Groups I, II, and III. DPOR and COR formed Group V (including chloroplast DPOR). The remaining Group IV consisted of only NFLs of unknown functions at the time this phylogenetic classification was proposed. Recently, two NFLs in Group IV were identified as CfbCD and MarHDK, revealing some of the functional diversity of NFLs [13–15]. Because most photosynthetic bacteria grow by photosynthesis under anaerobic conditions, they possess multiple functionally unknown NFLs in addition to nitrogenases and DPOR/COR. In the photosynthetic bacterium *Rhodobacter capsulatus*, COR was identified as the second NFL after DPOR [9, 10, 12]. Another set of NFL genes remains functionally unknown (Group IV) in the genome of *R. capsulatus*, which encodes a third NFL. Radical SAM enzymes (RSEs) catalyze a variety of reactions (FeMo-co biosynthesis, heme and (bacterio)chlorophyll biosynthesis, nucleotide biosynthesis, anaerobic pyruvate metabolism, etc.) via radical formation by electron transfer through intramolecular iron-sulfur clusters [23, 24]. Similar to nitrogenase, RSEs also require an anaerobic environment for their activities because the iron-sulfur clusters of RSEs are extremely vulnerable to oxygen [24]. Because NFLs and RSEs require anaerobic conditions in common, there are several pathways in which NFLs and RSEs act together or form a sequential reaction [3, 15, 25]. During the biosynthesis of FeMo-co, an RSE NifB produces the FeMo-co precursor NifB-co (L-cluster) [26], and an NFL NifEN functions as a scaffold for FeMo-co formation from NifB-co [3, 27]. In bacteriochlorophyll biosynthesis, an RSE BchE catalyzes the formation of protochlorophyllide followed by a sequential reaction of two NFLs, DPOR and COR [25, 28]. In addition, although their specific functions are unknown, an RSE MarB and an NFL MarHDK are essential components for methylthioalkane reduction [15]. Many RSEs with unknown functions remain in various prokaryote genomes, and some of them may function in collaboration with NFLs. Elucidation of the functions of these RSEs and NFLs not only reveals the unknown metabolic pathways of the organisms concerned but also provides a valuable clue to understanding the ancient metabolism that evolved under anaerobic environment at the dawn of life.

Sulfonate metabolism in prokaryotes plays a crucial role in sulfur cycles in various environments such as microbial mats and animal guts [29–31]. One of the sulfonates, taurine (aminoethylsulfonate), is an organic compound carrying amino and sulfo groups and is produced as a metabolite in the animal intestine [30–32]. A taurine-related compound, isethionate (hydroxyethylsulfonate), which has a hydroxy group instead of the amino group in taurine (Fig. 1), is not only a metabolic intermediate of taurine but also a sulfur compound that is naturally abundant in marine organisms [30, 32]. Taurine and isethionate are degraded to release sulfite, which is used as the sulfur source and as the terminal electron acceptor in anaerobic respiration [30]. Four metabolic pathways of sulfite formation from taurine and isethionate are known in prokaryotes (Fig. 1) [33–37]. The first pathway is catalyzed by sulfoacetaldehyde acetyltransferase (Xsc) [34]. Sulfoacetaldehyde is produced by the deamination of taurine (taurine pyruvate aminotransferase; Tpa or TauXY) [38, 39] or the oxidation of isethionate (isethionate dehydrogenase; IDH or IseJ or IseD) [39–41]. The second pathway is the conversion of isethionate to sulfite and acetaldehyde by isethionate lyase IslAB, which is an oxygen-independent reaction [35, 36]. The third pathway is the action of the NADH-dependent monooxygenase SsuED, which produces sulfite and glycolaldehyde from isethionate in an oxygen-dependent manner [33]. The fourth pathway is the oxygen-dependent conversion of taurine to sulfite by TauD [37]. *R. capsulatus* possesses Tpa and Xsc, allowing the production of sulfite from taurine to use the released sulfite as a sulfur source [42]. However, even in mutants in which both genes for Tpa and Xsc were disrupted, the ability to metabolize taurine was not completely lost [42]. Furthermore, no proteins showing significant similarity to IslAB, SsuED and TauD were found in the *R. capsulatus* genome. This suggests the existence of a novel sulfonate metabolic pathway in *R. capsulatus* that remains unknown to date.

**Fig. 1.**
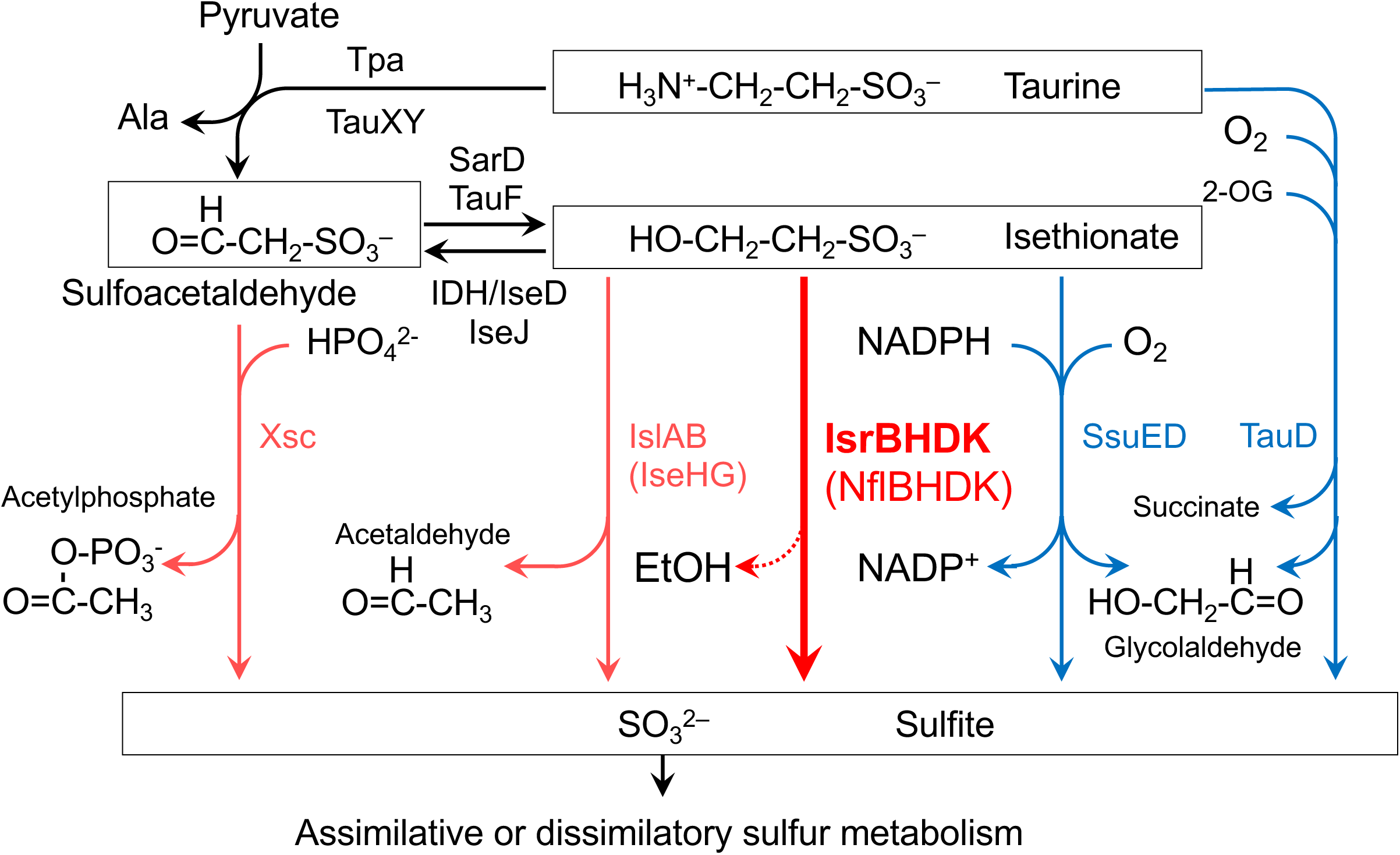
Metabolic pathways of taurine and isethionate in various bacteria. Oxygen-dependent and oxygen-independent desulfurization reactions are shown in blue and red, respectively. The new isethionate metabolic pathway (isethionate reductase; IsrBHDK) identified in this study is shown with the thick red arrow and in bold. Isethionate reductase releases sulfite from isethionate. However, the carbon product ethanol (EtOH) has not yet experimentally identified (dotted line).

Here we report that a third NFL in *R. capsulatus* is involved in a novel pathway for the use of isethionate as a sulfur source in anaerobic photosynthetic growth, and an RSE NflB is also essential for the pathway. The four genes for NFL and RSE form a tight gene cluster. The possible involvement of NFL and RSE in sulfonate metabolism was inferred from the known sulfonate metabolic gene existing at loci in the vicinity of this gene cluster. Analyses of the phenotype of targeted mutants, transcriptome, and genome profiling suggested that NFL and RSE are involved in a novel anaerobic isethionate metabolic pathway. This hypothesis was confirmed by the heterologous expression of the four genes for NFL/RSE and a putative isethionate transporter in a closely related species, *R. sphaeroides*, resulting in the ability to use isethionate as a sulfur source in this organism. This metabolic pathway, which is widely distributed among various prokaryotic phyla, plays an important role in sulfur metabolism in anaerobic environments. This discovery not only shows a new example of the cooperative actions of NFL and RSE but also provides a new perspective on sulfur metabolism at the dawn of life, which evolved under anaerobic environments.

## Results

The three genes, rcc02236, rcc02235, and rcc02234 that we focus on in this study encode functionally unknown NFL proteins that show significant similarity to the nitrogenase three subunits NifH, NifD, and NifK (44.1%, 16.8% and 16.4% from *Azotobacter vinelandii*, respectively), forming an operon similar to *nifHDK*. We tentatively call them *nflH*, *nflD*, and *nflK*, respectively (Fig. 2, Figs. S1-S3). In addition, a gene (rcc02237) immediately upstream of *nflH* encodes a protein that shows features of an RSE. We call this gene *nflB* after *nifB* (13.8% similarity to NifB of *A. vinelandii*; Fig. S4), which encodes an RSE involved in FeMo-co biosynthesis in the nitrogenase system. In the genome of *R. capsulatus*, genes for known sulfonate metabolism, such as *xsc* and *tpa*, are encoded in the vicinity of the *nflBHDK* gene cluster, implying that the proteins encoded by *nflBHDK* may also be involved in sulfonate metabolism (Fig. 2) [43].

**Fig. 2.**
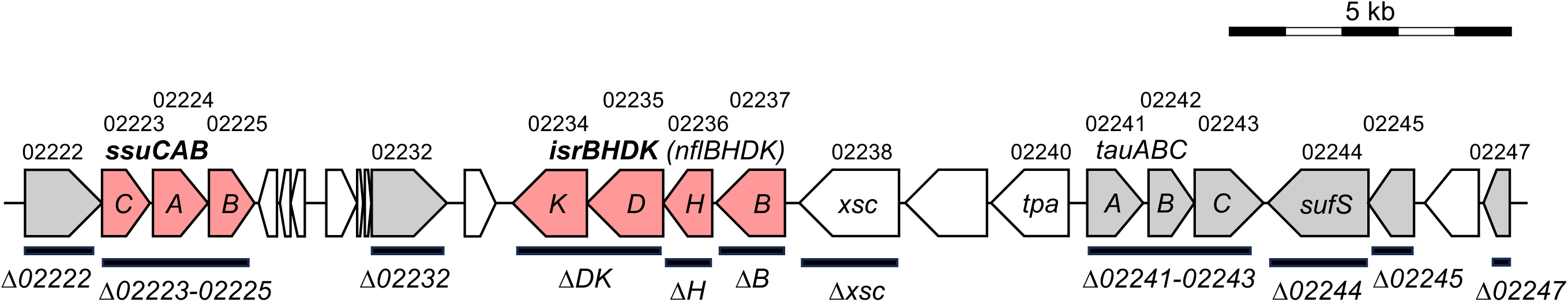
Gene arrangement of the 26-kb locus containing *isrBHDK* of the *R. capsulatus* genome. Genes shown by pink and gray are conserved specifically in the species carrying the *isrBHDK (nflBHDK)* genes in the genome profiling analysis (Table S3) and show high values (>100) of Ise/Sul ratio (Table 1). Genes with pink are the minimal set that conferred isethionate-dependent growth ability to *R. sphaeroides* (Table 2). The thick horizontal bars indicate the regions that were deleted in the targeted mutants (Fig. 3 and Table 2). Names of mutants are shown below the thick horizontal bars. The five-digit numbers on the gene map indicate the numbers following “rcc” of the locus tag.

**Table. 1.** List of genes specifically induced on medium with isethionate as the sole sulfur source (>100 Ise/Sul ratio) under anaerobic photosynthetic heterotrophic conditions.

| Locus tag<br>(rcc)<sup>a)</sup> | Annotation<sup>b)</sup> | Ise/Sul Ratio<sup>c)</sup> | FDR<sup>d)</sup> |
| --- | --- | --- | --- |
| 02113 | ornithine cyclodeaminase-1 | 129.0 | 3.04E-70 |
| 02143 | hypothetical protein | 4011.0 | 2.12E-73 |
| 02222 | ntaA; nitrilotriacetate monooxygenase, component A | 4968.0 | 1.62E-83 |
| 02223 | ABC transporter, permease protein (SsuC) | 4662.3 | 7.74E-53 |
| 02224 | ABC transporter, substrate-binding protein (SsuA) | 3422.2 | 6.14E-130 |
| 02225 | ABC transporter, ATP-binding protein (SsuB) | 1259.4 | 1.26E-25 |
| 02226 | conserved hypothetical protein | 795.1 | 3.93E-44 |
| 02227 | conserved hypothetical protein | 1148.0 | 6.71E-24 |
| 02228 | conserved hypothetical protein | 2810.0 | 2.53E-48 |
| 02229 | protein of unknown function DUF1234 | 2816.7 | 1.29E-119 |
| 02230 | membrane protein, putative | 423.8 | 7.88E-20 |
| 02231 | hypothetical protein | 407.9 | 1.16E-19 |
| 02232 | FAD dependent oxidoreductase | 3987.9 | 1.66E-117 |
| 02233 | conserved hypothetical protein | 1386.9 | 1.19E-52 |
| 02234 | oxidoreductase/nitrogenase, component 1 (IsrK) | 1748.9 | 5.70E-64 |
| 02235 | oxidoreductase/nitrogenase, component 1 (IsrD) | 1148.9 | 4.24E-88 |
| 02236 | nifH2; nitrogenase iron protein-2 (IsrH) | 1819.7 | 8.72E-65 |
| 02237 | radical SAM family protein (IsrB) | 4787.6 | 3.07E-76 |
| 02241 | tauA; taurine ABC transporter, periplasmic taurine-binding protein TauA | 931.6 | 1.56E-48 |
| 02242 | tauB; taurine ABC transporter, ATP-binding protein TauB | 488.4 | 2.52E-30 |
| 02243 | tauC; taurine ABC transporter, permease protein TauC | 687.1 | 5.85E-26 |
| 02244 | sufS2; cysteine desulfurase-2 | 1421.7 | 6.32E-110 |
| 02245 | major membrane protein I | 1292.6 | 1.06E-149 |
| 02246 | serine O-acetyltransferase-2 | 1215.4 | 1.78E-84 |
| 02247 | rhodanese domain protein | 1005.3 | 5.27E-37 |
| 02281 | dimethyl sulfoxide reductase, A subunit | 264.3 | 2.20E-16 |
| 02535 | sulfate ABC transporter, permease protein CysW | 182.8 | 3.24E-43 |
| 02536 | sulfate ABC transporter, permease protein CysT | 289.4 | 6.28E-46 |
| 02647 | bifunctional protein PutA | 116.1 | 6.42E-168 |
| 02744 | sulfate/thiosulfate ABC transporter, periplasmic<br>sulfate/thiosulfate-binding protein | 4603.8 | 4.86E-260 |

<sup>a)</sup> The five-digit number indicates the number following the locus tag of rcc
<sup>b)</sup> Annotations in GenBank data base. The names of proteins proposed in this work are shown in parentheses.
<sup>c)</sup> The ratio of transcripts levels of the gene in cells grown in media containing isethionate (Ise) and sulfate (Sul) as the sole sulfur sources, respectively.
<sup>d)</sup> FDR calculated by DEseq2

**Table. 2.**
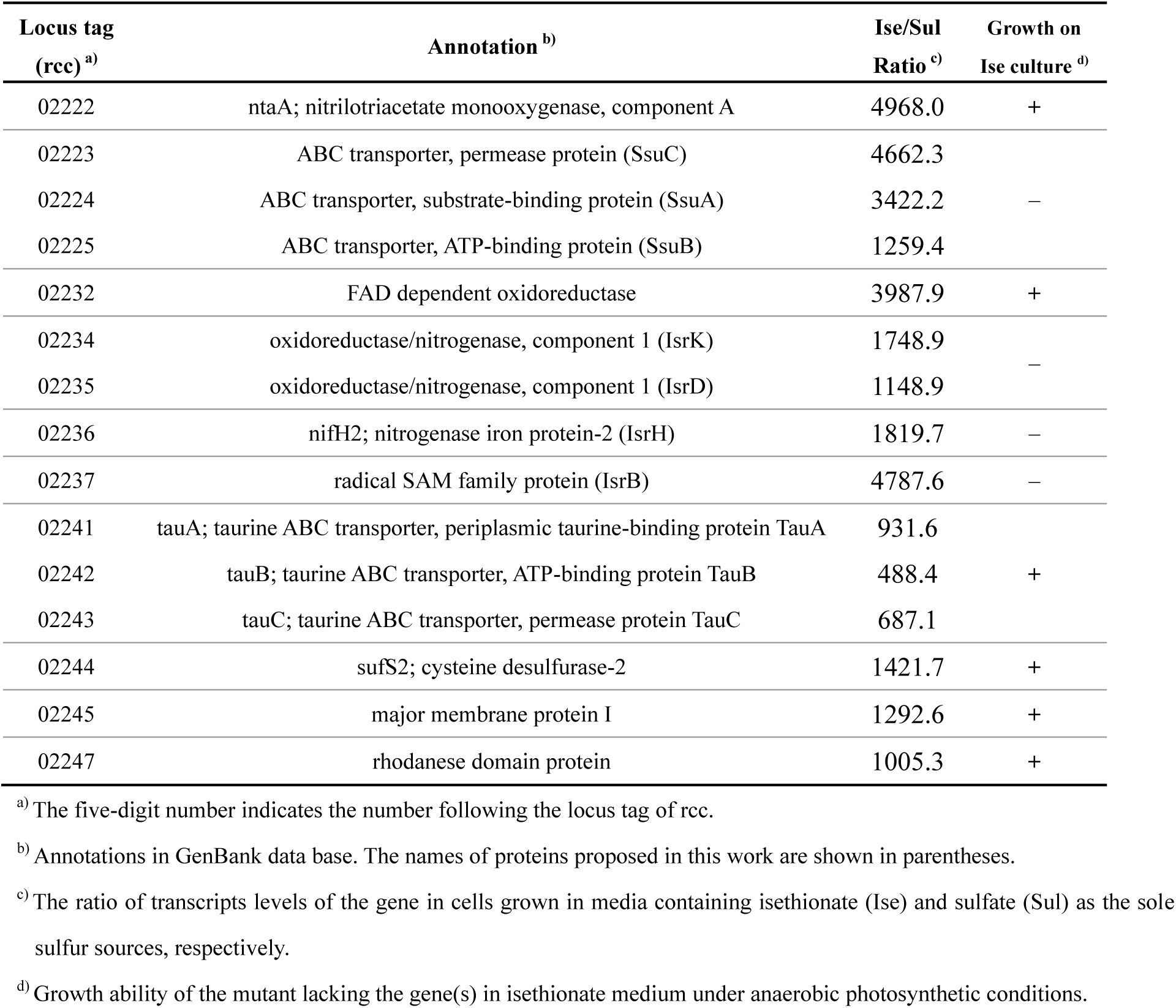
List of genes conserved specifically in *R. capsulatus* identified by genome profiling analysis and specifically induced on medium with isethionate as the sole sulfur source (>100 Ise/Sul ratio) and the growth ability of the targeted disrupted mutants under anaerobic photosynthetic heterotrophic conditions on isethionate medium.

To elucidate the function of these genes in terms of sulfonate metabolism, we first examined photosynthetic growth of *R. capsulatus* with two sulfonates, taurine and isethionate, as the sole sulfur sources (Fig. 3). *R. capsulatus* WT (see Materials and Methods) grew well with both sulfonates and sulfate, indicating that *R. capsulatus* can utilize both taurine and isethionate as the sole sulfur sources. To confirm the metabolic pathway of these sulfonates in *R. capsulatus*, we isolated an *xsc*-knockout mutant, *Δxsc*, and examined photosynthetic growth (Fig. 3). In contrast to the loss of growth ability of Δ*xsc* in taurine medium, *Δxsc* grew well with isethionate as the sole sulfur source, indicating that Xsc is essential for taurine assimilation but not for isethionate. Given that *R. capsulatus* does not have genes corresponding to the *islAB* and *ssuED* genes (Fig. 1), this result suggested the presence of a novel alternative *xsc*-independent metabolic pathway for isethionate in *R. capsulatus* under anaerobic photosynthetic conditions.

**Fig. 3.**
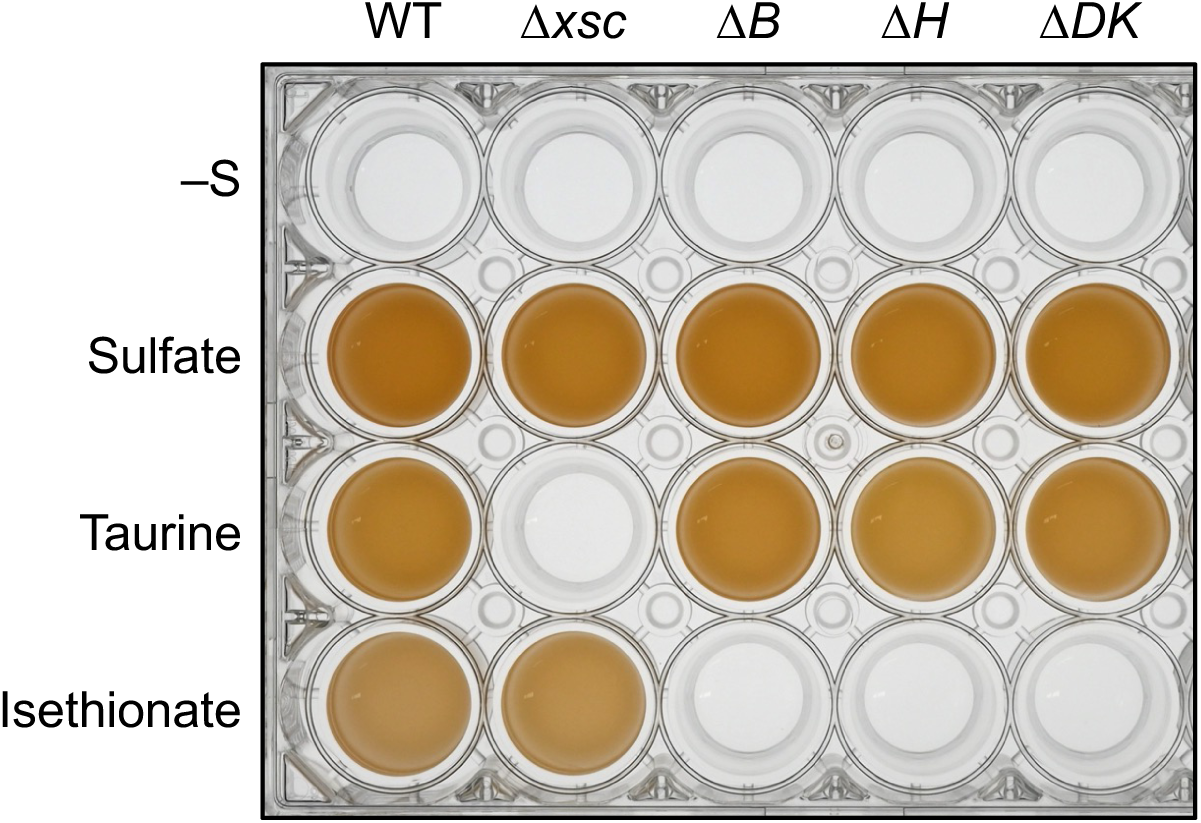
Growth of WT and four targeted mutants under anaerobic photosynthetic conditions with three sulfur compounds. WT and the four mutants were grown with various sulfur sources in screw-capped test tubes under anaerobic photosynthetic conditions. The cultures without sulfur (–S), with sulfate, taurine, and isethionate were incubated for 120 h, 48 h, 72 h, and 120 h, respectively. The resultant cultures were transferred into a 24-well plate to show clearly their growth.

Next, to obtain clues to genes involved in the novel isethionate metabolic pathway in *R. capsulatus*, transcriptome analysis (RNA-seq) was performed to globally extract genes that are specifically induced under growth conditions to use isethionate as a sulfur source. We assumed that genes involved in isethionate metabolism are transcriptionally repressed when growing with sulfate as the sulfur source and induced when growing with isethionate as the sole sulfur source. *R. capsulatus* WT was cultivated photosynthetically in a medium containing sulfate or isethionate as the sole sulfur source. The ratios of transcript levels grown with isethionate to those grown with sulfate (the Ise/Sul ratio) were estimated for all genes. Thirty genes were extracted with Ise/Sul ratios greater than 100 (Table 1), 23 of which were located at the chromosomal locus spanning approximately 26 kb from rcc02222 to rcc02247 (Fig. 2, Fig. S5). The *nflBHDK* gene cluster was contained this chromosomal locus and showed high Ise/Sul ratios (1,300 to 5,000). In addition, other genes in the 26-kb locus, such as the *tauABC* genes for the ABC-type taurine transporter (rcc02241–rcc02243), genes for the ABC-type sulfonate transporter (rcc02223–rcc02225), and *sufS* for cysteine desulfurase (rcc02244), also showed high Ise/Sul ratios. Four genes, rcc02222 (monooxygenase-type enzyme), rcc02228, rcc02229, and rcc02232 (FAD dependent oxidoreductase), whose functions are unknown, also showed Ise/Sul ratios of more than 2,000 (Table 1). In contrast, the Ise/Sul ratios of *xsc* and *tpa* were less than 100, even though these genes are located at just upstream of *nflBHDK* (Fig. S5). The RNA-seq results indicate that *nflBHDK* is involved in a novel isethionate metabolic pathway.

Then, targeted mutants for *nflB*, *nflH*, and *nflDK* (*ΔB*, *ΔH*, and *ΔDK*, respectively) were isolated (Fig. 2) and photosynthetic growth was examined in isethionate medium (Fig. 3). All three mutants lost the ability to grow in the isethionate medium. These results indicated that all four *nflBHDK* genes are essential for growth using isethionate as the sole sulfur source.

To specify the reaction involving NflBHDK, an expression plasmid of the *islAB* genes encoding isethionate lyase that catalyzes the degradation of isethionate to sulfite and acetaldehyde from the enterobacterium *Bilophila wadsworthia* 3.1.6 was introduced into *ΔDK* strain. The isolated transconjugant *ΔDK+islAB* recovered growth in isethionate medium (Fig. 4). This result indicates that the isethionate lyase reaction can substitute for a reaction involving NflDK to support isethionate-dependent growth. Given that sulfite is converted to sulfide by sulfite reductase in *R. capsulatus* to use as a sulfur source, this result indicates that the reaction catalyzed by NflDK, which IslAB can replace, contains at least the release of sulfite from isethionate. Furthermore, considering that NFLs catalyze a variety of reductions, the *nflHDK* genes are most likely to encode subunits of a novel enzyme, isethionate reductase, that catalyzes the reductive cleavage the C-S bond of isethionate. On the basis of these results, we propose to rename *nflBHDK* as *isrBHDK* after <u>is</u>ethionate reductase.

**Fig. 4.**
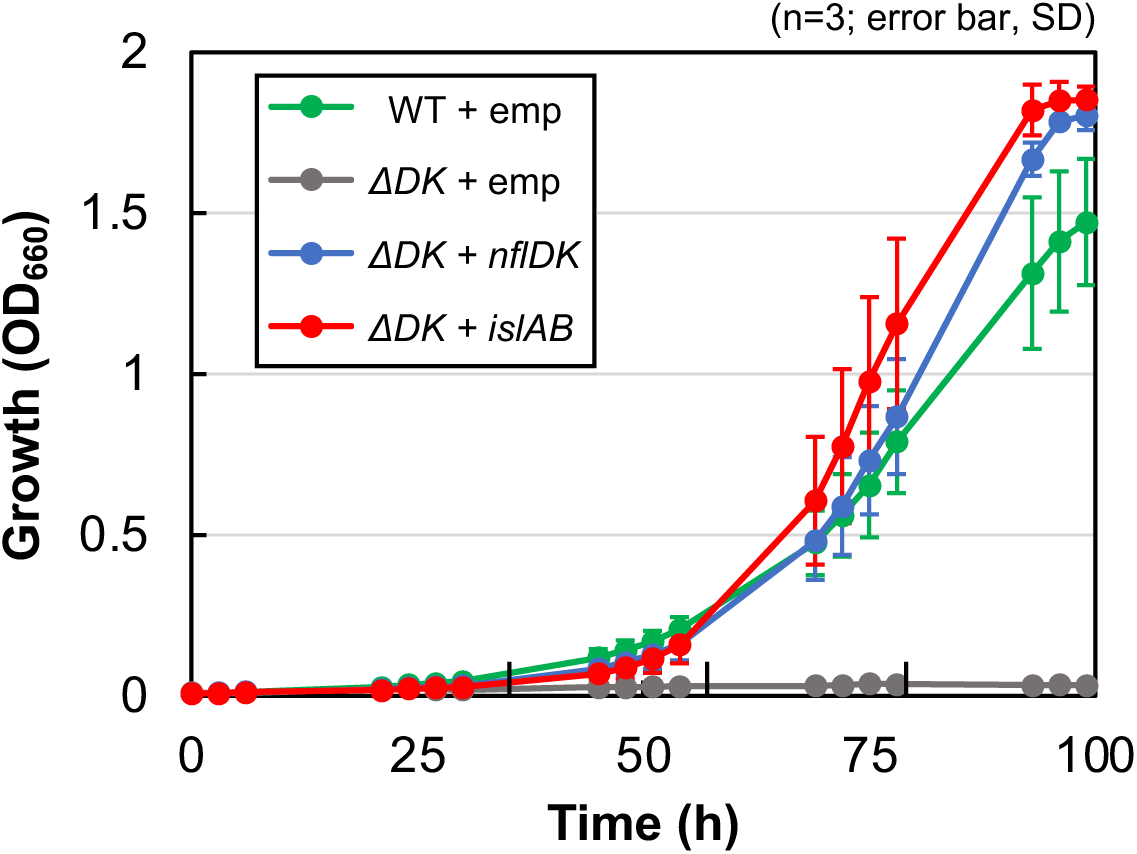
Complementation of growth ability of *ΔDK* in isethionate medium with expression of *islAB*. Transconjugants carrying the empty vector (WT/Δ*DK* + emp), and *ΔDK* carrying *nflDK* (Δ*DK+nflDK*) and *islAB* (Δ*DK*+*islAB*) were grown in isethionate media under anaerobic photosynthetic conditions. Growth was monitored with optical density at 660 nm (OD_660_).

To clarify whether the four genes *isrBHDK* are sufficient for the availability of isethionate as a sulfur source or other genes are required, genome profiling analysis was performed. In this *in silico* analysis, genes specifically conserved in organisms carrying *isrBHDK* are extracted by genome comparison among closely related species. Eight species, including *R. capsulatus* SB1003 (*R. capsulatus*_B in the GTDB classification), were selected as a group in which *isrBHDK* genes are conserved. Five species including *R. capsulatus* DSM1710 and 17 species including *R. sphaeroides* 2.4.1 (*R. capsulatus* and *Celeibacter*_A *sphaeroides* in the GTDB classification, respectively) were selected as the other group in which *isrBHDK* genes are missing (Fig. S7, Tables S1, S2). Using the genome information of these 30 species, 121 genes were extracted as conserved only in the eight species carrying *isrBHDK* genes (Table S3). Interestingly, 11 of the 121 genes were located in the 26-kb gene cluster containing *isrBHDK*, with an Ise/Sul ratio of > 100 in *R. capsulatus* (Fig. 2). We then regarded the proteins encoded by the 11 genes as strong candidates acting together with IsrBHDK, and targeted mutants were isolated for all 11 genes. For the two sets of genes for the ABC-type transporter in the 11 genes, a single targeted mutant with all three genes removed was isolated, respectively, rather than three targeted mutants lacking individual genes. Then, the seven targeted mutants were isolated and examined for photosynthetic growth in isethionate medium (Fig. 2, S6). Only one mutant Δrcc02223-rcc02225 lacking the three genes (rcc02223, rcc02224, and rcc02225) encoding subunits of the ABC-type sulfonate transporter failed to grow in isethionate medium other than the three mutants for *isrBHDK* (Fig. S6, Table 2). This result indicates that the three genes for the putative isethionate transporter are essential for growth in the isethionate medium and that the transporter acts with IsrBHDK to support photosynthetic growth using isethionate as the sole sulfur source. Because the three subunits of the ABC-type sulfonate transporter (rcc02223, rcc02224 and rcc02225) show high similarity to those of subunits of the ABC-type sulfonate transporter SsuC, SsuA, and SsuB from *E. coli* with sequence similarity of 39.2%, 28.5%, and 45.7%, respectively, we propose to designate these three genes, rcc02223, rcc02224, and rcc02225, as *ssuC*, *ssuA*, and *ssuB*, respectively, in *R. capsulatus* SB1003 [44].

Genome profiling and reverse genetic analysis suggest that IsrBHDK and SsuCAB are sufficient to use isethionate as the sole sulfur source. Next, we examined whether heterologous expression of these seven genes confers the ability to use isethionate as the sole sulfur source to a different host organism. *R. sphaeroides* 2.4.1, which is a close relative of *R. capsulatus* but lacks the *isrBHDK* genes and is unable to grow in media containing isethionate as the sole sulfur source (Fig. S8). A shuttle vector expressing *isrBHDK* and *ssuCAB* of *R. capsulatus* was introduced into *R. sphaeroides*, and the transconjugant IsrBHDK+SsuCAB was isolated. IsrBHDK+SsuCAB showed significantly enhanced growth in isethionate medium compared with the control transconjugant carrying the empty vector (Fig. 5). Furthermore, such growth enhancement was not observed in transconjugant carrying a plasmid expressing only either *isrBHDK* or *ssuCAB*. The results indicate that isethionate in the medium is taken up by the SsuCAB transporter, IsrBHDK releases sulfite from isethionate, and sulfite is used as the sulfur source. It was also strongly suggested that the four genes *isrBHDK* are the minimum gene set required for the novel isethionate assimilatory enzyme Isr to function.

**Fig. 5.**
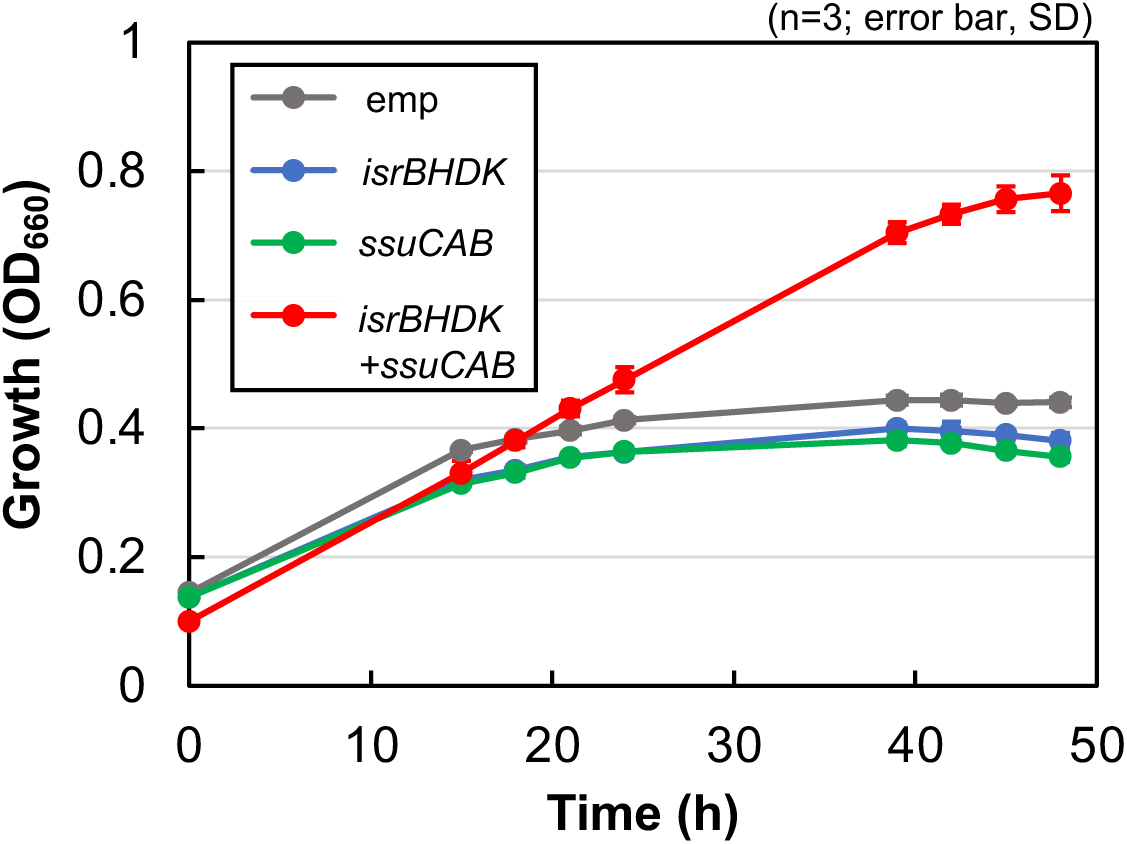
Expression of *isrBHDK* and *ssuCAB* conferred of isethionate assimilation ability to *R. sphaeroides*. Plasmids expressing *isrBHDK*, *ssuCAB* (rcc02223-02225), and both gene clusters of *isrBHDK* and *ssuCAB* (*isrBHDK+ssuCAB)* were introduced into *R. sphaeroides* by conjugation, and growth of the transconjugants were monitored by OD_660_ in culture medium containing isethionate as the sole sulfur source under anaerobic photosynthetic conditions. The transconjugant carrying only the empty vector (emp) was the negative control.

## Discussions

In this study, we demonstrate that the NFL (IsrHDK) and the RSE (IsrB) in *R. capsulatus* potentially function as the novel enzyme isethionate reductase Isr under anaerobic conditions. IsrHDK is a third NFL whose function was identified after COR in *R. capsulatus*. In addition, we identified the ABC-type isethionate transporter SsuCAB in *R. capsulatus*. The cooperative actions of IsrBHDK and SsuCAB allow *R. capsulatus* to use isethionate in the environment as a sulfur source (Fig. 6).

**Fig. 6.**
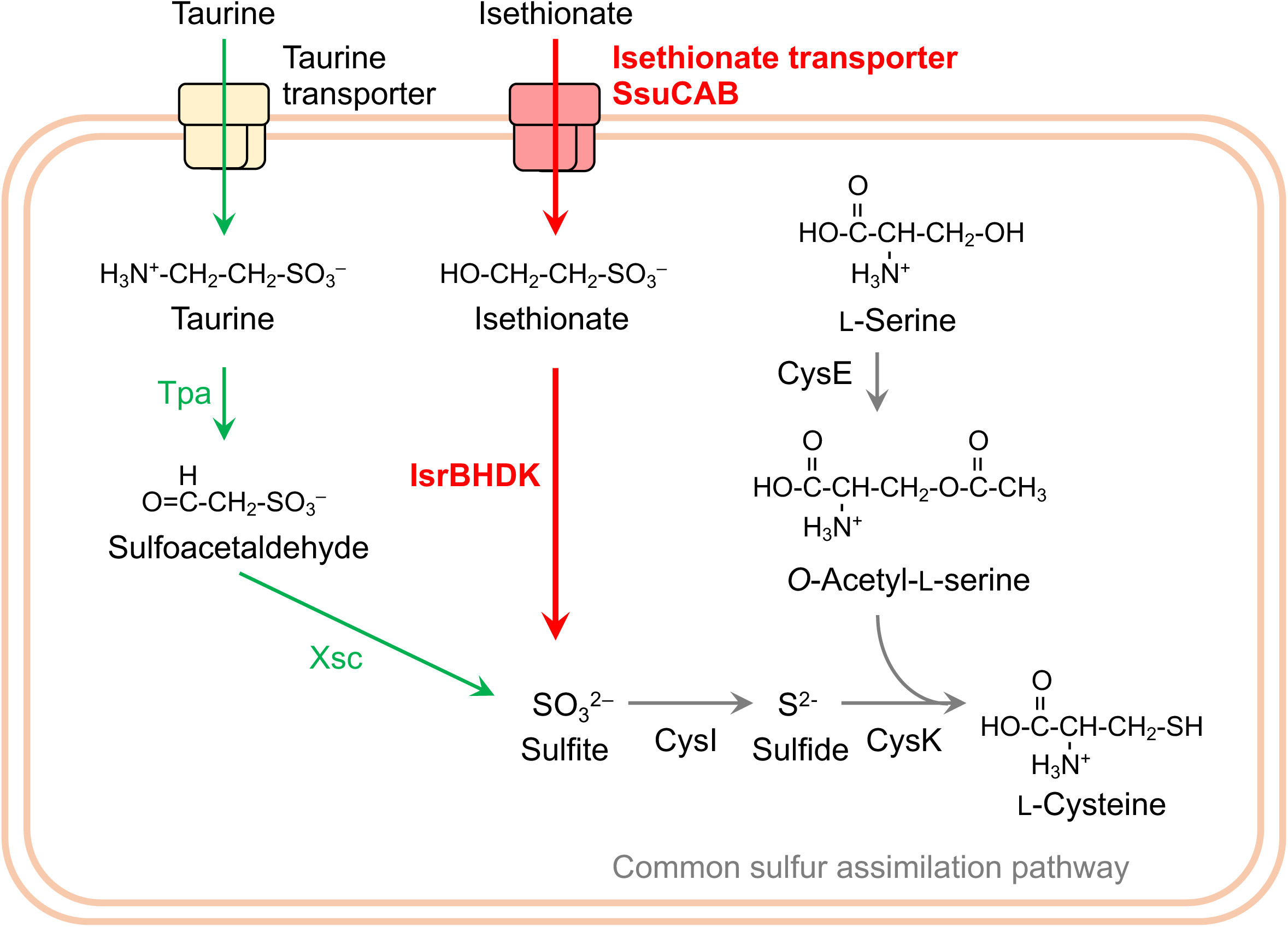
Metabolic pathway of isethionate and taurine in *R. capsulatus* SB1003. Metabolic pathways specific for isethionate and taurine are shown in red and green, respectively. Isethionate reductase (IsrBHDK) and isethionate transporter (SsuCAB) identified in this study are shown in bold. SsuCAB uptakes isethionate in the medium to the cytoplasm and isethionate reductase degrades isethionate to release sulfite, followed by the sulfur assimilation pathways common to many bacteria indicated by gray arrows.

### Cysteine conservation in the amino acid sequences of IsrHDK

Based on the structural features of nitrogenases and NFLs, DPOR and COR, which have been biochemically studied, Isr may consist of two components: a reductase component (a IsrH dimer) and a catalytic component (a IsrDK heterotetramer) [9–12, 19, 21]. The conservation of Cys residues, which serve as ligands for iron-sulfur clusters in various iron-sulfur proteins, provides an important clue for predicting the structure and function of metal clusters retained in NFLs. The NifH homolog IsrH has two fully conserved Cys residues (Cys99 and Cys135) that hold a [4Fe-4S] cluster in the nitrogenase Fe protein (NifH), in addition to the full conservation of the ATP-binding motif (Fig. S1) [3]. Thus, IsrH, like other NifH homologs, is expected to hold a single [4Fe-4S] cluster in a homodimer and functions as a reductase component that transfers electrons to the catalytic component IsrDK, which is coupled with ATP hydrolysis [3, 18, 45].

IsrD, the NifD homolog, has only a single conserved Cys74 corresponding to Cys88 of the three Cys residues (Cys62, Cys88, and Cys154) in NifD of *A. vinelandii* involved in P-cluster chelation in the MoFe protein (Fig. S2) [3]. The other two residues corresponding to Cys62 and Cys154 are conserved as Arg and Pro/Ser in IsrD. In contrast, all other NifD homologs (CfbD, MarD, NfaD [46], BchN/ChlN, and BchY) have three conserved Cys in the corresponding region. Regarding the NifK homolog, IsrK, the three Cys (Cys70, Cys95, and Cys153 in NifK of *A. vinelandii*), which are involved in P-cluster chelation, were completely conserved as Cys12, Cys37, and Cys99 (Fig. S3). The conservation of Cys residues in IsrD and IsrK (one and three Cys residues in IsrD and IsrK, respectively) suggests that these four Cys may be involved in the holding of metal clusters such as a [4Fe-4S] cluster. It is noteworthy that in the NB protein of DPOR (BchN-BchB), one [4Fe-4S] cluster (the NB cluster) is held by three Cys in BchN and one Asp in BchB [11, 47]. Thus, it cannot be excluded that residues other than Cys may be involved in metal cluster chelation.

In addition, neither Cys275 nor His442 of the NifD subunit (*A. vinelandii*) involved in chelating FeMo-co is conserved in IsrD, suggesting that it is unlikely to hold a complex metal cluster such as FeMo-co. This is a similar feature of BchN/ChlN (DPOR) and BchY (COR) in Group V NFLs (Fig. S2). In DPOR, the substrate protochlorophyllide binds to the site in the NB protein that largely corresponds to the site occupied by FeMo-co in the MoFe protein [3, 11]. Given that NFLs share common structural features, IsrDK may also bind the substrate isethionate in a site corresponding to the FeMo-co binding site of the MoFe protein similar to the NB protein.

### Reaction catalyzed by IsrBHDK

The biochemical reactions catalyzed by known NFLs implied that in general, NFLs catalyze a variety of reductions, in which the catalytic component (homolog of DK subunit) reduces diverse substrates using electrons from the reductase component (homolog of H subunit) [4, 5]. In addition to this implication, given that the expression of IslAB complemented the ability to use isethionate as the sulfur source in the mutant *ΔDK*, it is suggested that Isr catalyzes the reductive cleavage of the C-S bond in isethionate releasing sulfite. This cleavage reaction appears to be similar to the reaction catalyzed by Mar with respect to the reductive cleavage of the C-S bond [15]. Mar converts 2-(methylthio)ethanol, giving rise to methane thiol, ethylene, and water. If it is a simple reductive cleavage of the C-S bond of isethionate by Isr, the resulting carbon product is presumed to be ethanol (Fig. 1).

### Function of IsrB

The loss of growth ability in isethionate medium in the targeted mutant of *isrB* (*ΔB*, Fig. 3) strongly indicated that IsrB is an essential protein for the reaction catalyzed by Isr. What is the function of IsrB in isethionate reduction? In the FeMo-co biosynthesis of the nitrogenase system, NifB is involved in the formation of NifB-co, the precursor of FeMo-co [3, 27]. In this process, NifB catalyzes a series of complex reactions to extract a methyl group from *S*-adenosylmethionine (SAM) as the first reaction, followed by the conversion of two [4Fe-4S] clusters to NifB-co with the insertion of the methyl carbon as the central carbide of NifB-co [3, 26, 48]. In the first reaction, NifB cleaves the C-S bond of SAM to liberate the methyl group. In this respect, it would be similar to the reductive cleavage of the isethionate C-S bond that we hypothesize that IsrHDK catalyzes. As another example, one subunit of isethionate lyase, IslB, is an RSE that acts as the activating the glycyl radical enzyme IslA to form a glycyl radical on the Gly residue of IslA [35, 36]. This function of the activating enzyme via radical formation is also observed in PflA, the RSE unit of pyruvate formate lyase (PflB) [49]. Currently, it is unclear whether IsrB cleaves the C-S bond or activates IsrHDK. Similarly, the role of MarB in the methylthioalkane reductase system is also unclear [15]. Further study from a biochemical perspective is required.

### Distribution of isethionate-assimilatory enzymes

Xsc and IslAB are assimilatory enzymes that catalyze the desulfurization reactions of sulfoacetaldehyde and isethionate, respectively, in the oxygen-independent manner (Fig. 1) [34–36]. In this study, we identified the genes for Isr as a third oxygen-independent desulfurizing enzyme. The distribution of these three enzymes; Xsc, Isl, and Isr, in prokaryotes shows that Isr is almost exclusively distributed in the phyla of *Bacillata*, *Spirochaeta*, *Pseudomonadota*, and *Myxococcota* with a few exceptions (Fig. 8). We predict that these organisms carrying Isr without Xsc and IslAB can degrade isethionate to sulfite for use as a sulfur source (assimilation) or as an electron acceptor (dissimilation) under anaerobic conditions. The distribution of the three enzymes suggests that most prokaryotes carry only one enzyme for isethionate desulfurization, and organisms with multiple enzymes such as Xsc and Isr, as in *R. capsulatus*, are rare.

*R. capsulatus* may use Xsc and Isr differentially for taurine and isethionate, respectively. In fact, in *R. capsulatus*, the ability to metabolize isethionate was not affected in *Δxsc*, and the ability to metabolize taurine was not affected in any of the *isr* mutants (*ΔB*, *ΔH*, and *ΔDK*). In addition, RNA-seq analysis showed that the Ise/Sul ratios of *xsc* and *tpa* expression levels were very low in contrast to more than 1,000 of the Ise/Sul ratios of *isrBHDK* expression (Fig. S5), indicating that isethionate addition does not significantly enhance the transcript levels of *xsc* and *tpa*. These results indicate that *xsc* and *tpa* are regulated by factors other than isethionate, while they are located just adjacent to *isrBHDK*, which is strongly induced by isethionate, supporting the idea that the metabolic pathways for isethionate and taurine are independent in *R. capsulatus* (Fig. 6).

### Molecular phylogenetic aspects of IsrBHDK

Molecular phylogenetic analysis of IsrBHDK was performed to understand the evolutionary relationship of this novel isethionate reductase with other NFLs and RSEs. First, a phylogenetic tree was constructed on the NifD homologs from various species. IsrD formed an independent clade in close proximity to MarD and NfaD within Group IV (Fig. 7A). A similar pattern was observed in the molecular phylogenetic trees of NifH homologs (Fig. 7B), suggesting that IsrH and IsrD originated from the common ancestral NFL sharing with MarHDK and NfaHDK within Group IV, but evolved to form independent clades. Interestingly, the phylogenetic analysis of the RSEs showed a completely different phylogenetic relationship among IsrB, MarB, NfaB, and NifB (Fig. 7C). In fact, the sequence similarity in these B subunits (IsrB, MarB, NfaB, and NifB) is limited to the radical SAM motif (Fig. S4). A blast search against the swissprot database revealed that the protein with the highest similarity to IsrB was BciD. BciD is involved in the conversion of the C7-methyl group to the C7-formyl group in bacteriochlorophyll *e* biosynthesis [50]. This phylogenetic analysis indicates that the phylogenetic relationship between IsrB and NifB is different from that between IsrHDK and NifHDK. In other words, the cooperative action of NFL and RSE in Isr could be created through the recruitment of genes from independent lineages in the early evolution of life.

**Fig. 7.**
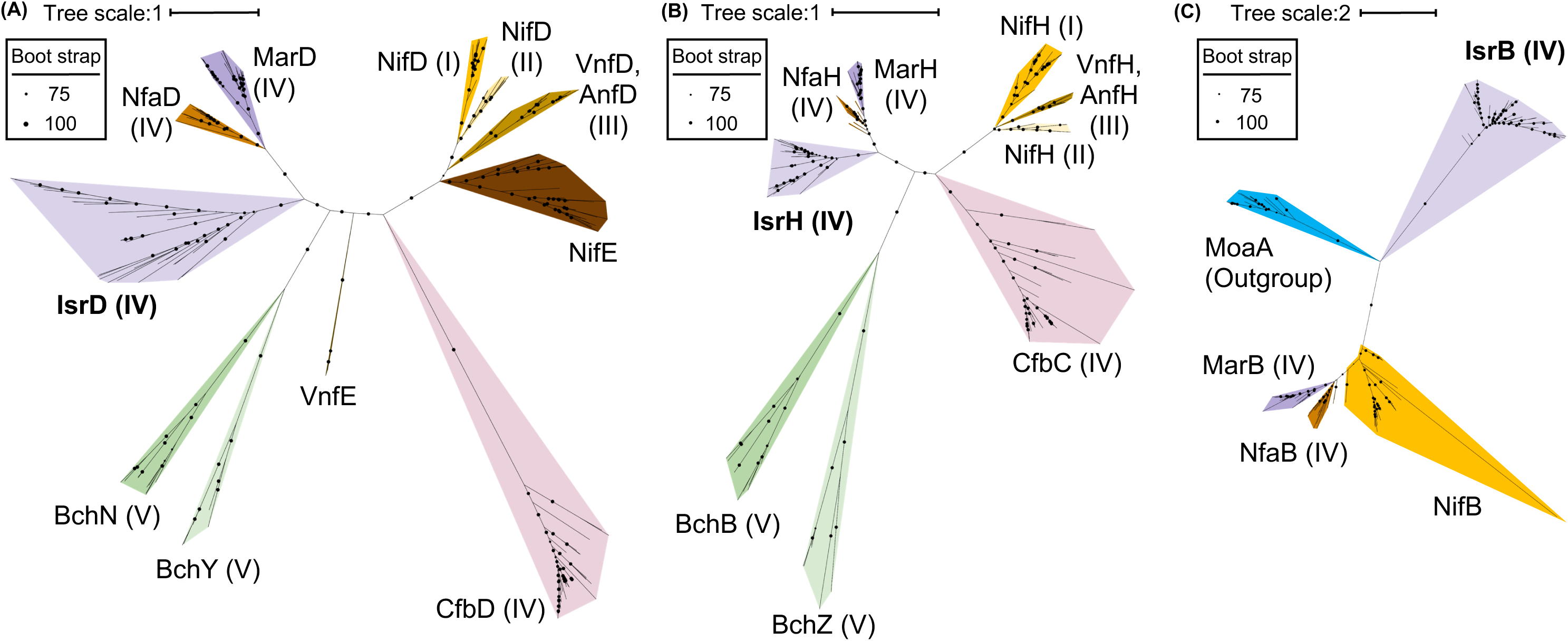
Phylogenetic trees of IsrD (A), IsrH (B), and IsrB (C) and their homologs. (**A**) Phylogenetic tree of IsrD and its homologs, (**B**) IsrH and its homologs, and, (**C**) IsrB and its homologs. Groups I–V proposed by Raymond et al. (2004), to which each protein belongs, are shown in parentheses. NifE and VnfE were not included in the phylogenetic tree in Raymond et al. (2004).

**Fig. 8.**
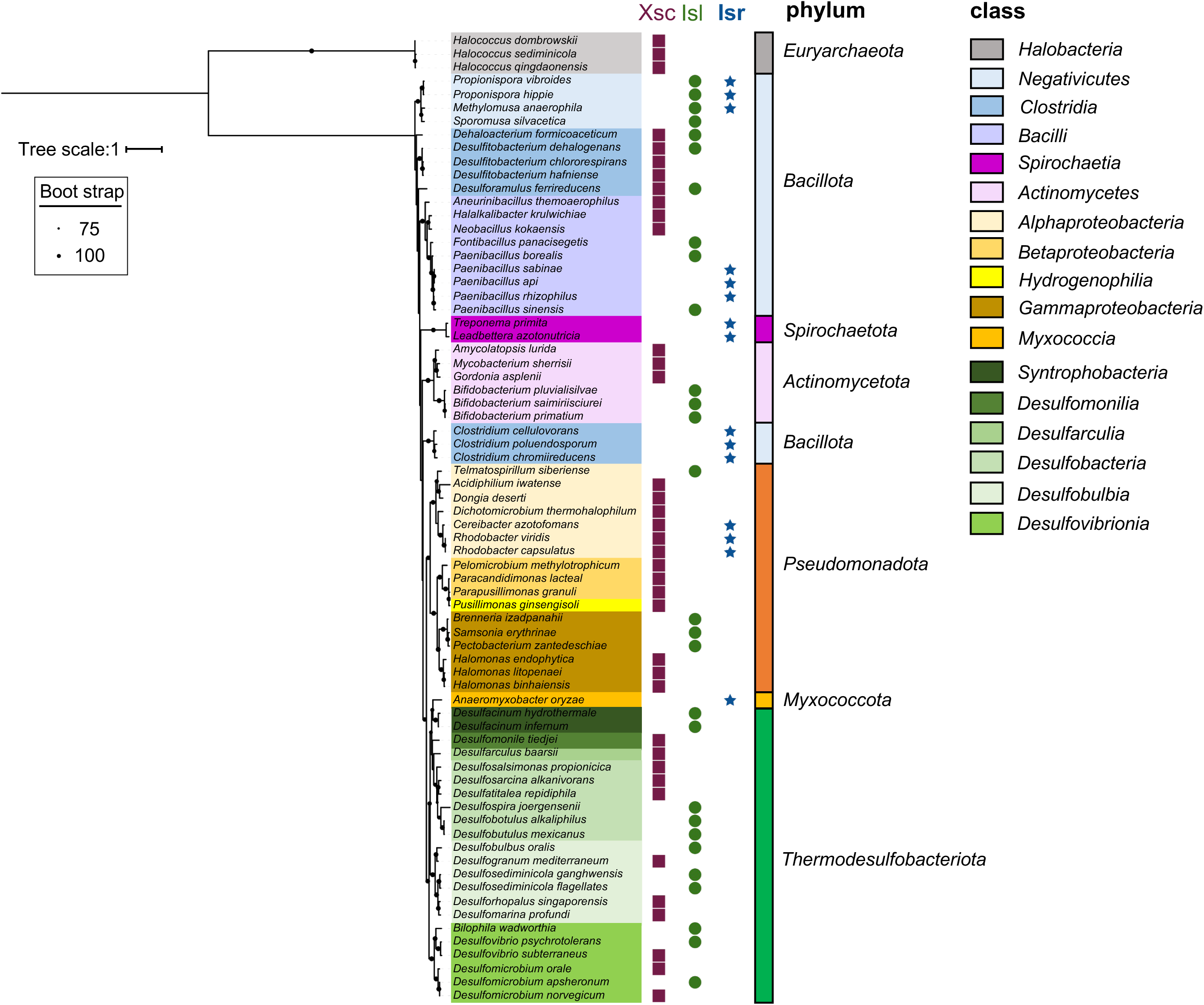
Distribution of the three oxygen-independent isethionate metabolic enzymes (Isr, Isl, and Xsc) in prokaryotes. Distribution of the three enzymes; Xsc (purple squares), IslA (green circles), and IsrD (blue stars), which are oxygen-independent enzymes for isethionate metabolism, are plotted on the phylogenetic tree of 16s rRNA.

### Aerobic isethionate metabolism

The *isr* mutants lost the ability to grow with isethionate as a sulfur source under anaerobic photosynthetic conditions, whereas the mutants were still able to use isethionate as a sulfur source under aerobic heterotrophic conditions (Fig. S9). The isethionate-dependent growth capacity under aerobic conditions was observed even in *Δxsc*, indicating that an aerobic isethionate metabolic pathway operates independently of the Xsc and Isr systems. SsuED is the enzyme responsible for oxygen-dependent isethionate metabolism under aerobic conditions in *E. coli*. However, no genes homologous to *ssuED* have been found in the *R. capsulatus* genome, implying that there is also a novel aerobic isethionate-metabolizing enzyme that has not been reported so far. Given that isethionate-dependent growth is completely lost under anaerobic conditions in the *isr* mutants, this aerobic enzyme (or pathway) should include oxygen-dependent reactions.

The coexistence of oxygen-independent (oxygen-labile) and oxygen-dependent (oxygen-tolerant) enzymes in isethionate metabolism is reminiscent of the coexistence of these enzymes in the heme and chlorophyll biosynthesis pathways. DPOR, the NFL responsible for protochlorophyllide reduction in chlorophyll biosynthesis, coexists with an oxygen-tolerant and light-dependent enzyme; protochlorophyllide oxidoreductase (LPOR) in cyanobacteria, algae, and lower plants [51, 52]. In addition, oxygen-labile anaerobic enzymes (HemN and BchE) and oxygen-dependent enzymes (HemF and AcsF) coexist in the eighth reaction (coproporphyrinogen oxidation; HemN and HemF) in heme biosynthesis [53–55] and in the reaction of protochlorophyllide formation (Mg-protoporphyrin IX monomethyl ester cyclase; BchE/ChlE and AcsF/ChlA) in the bacteriochlorophyll and chlorophyll biosynthesis systems [25, 56, 57]. In these steps, the oxygen-labile anaerobic enzymes (HemN and BchE) are RSEs. The coexistence of the newly identified anaerobic enzyme Isr and the presumed aerobic enzyme can be regarded as a new example of adaptive evolution to aerobic environments that emerged after the Great Oxidation Event (GOE) in the early evolution of life [58–60]. Elucidation of the functions of NFLs and RSEs with unknown functions is important not only for the metabolic capacity of the organisms concerned but also for elucidating ancestral metabolism and following adaptive evolution in response to GOE.

## Materials and Methods

### 1. Strains and culture conditions

In this study, an adaptive strain that grows without a lag time in RCV-isethionate medium [61] was isolated from a single colony of *R. capsulatus* SB1003 on an RCV-isethionate medium agar plate and used as the wild type (WT). Both WT and gene disrupted mutants were pre-cultured in 3-mL PY medium overnight and then inoculated into various media (initial OD_660_ = 0.01): RCV medium without any sulfur compounds (RCV(-S)), with MgSO_4_ (RCV-SUL), taurine (RCV-TAU), and sodium isethionate (RCV-ISE). The final concentrations of MgSO_4_, taurine, and isethionate in the RCV-SUL, RCV-TAU, and RCV-ISE media were 0.1 mM, respectively. These RCV media were prepared by substituting (NH_4_)_2_SO_4_, MgSO_4_, and FeSO_4_ with NH_4_Cl, MgCl_2_, and FeCl_3_, respectively. For anaerobic photosynthetic growth, liquid cultures were prepared in 30-ml test tubes sealed with air-tight screw caps and illuminated with a 75-W incandescent bulb (National, Osaka, Japan) at 34°C.

*R. sphaeroides* J001-1, an adapted strain for growth in Sistrom medium [62] derived from the rifampicin-resistant strain *R. sphaeroides* J001, was used in this study [63]. To confirm its ability to use isethionate, *R. sphaeroides* J001-1 was cultured under anaerobic photosynthetic conditions in PYS [64] medium for 48 h as a preculture. Precultured cells were washed with Sistrom (-S) medium and inoculated into Sistrom (-S) medium containing 0.1 mM isethionate. Sistrom (-S) was prepared by substituting (NH_4_)_2_SO_4_, MgSO_4_, and FeSO_4_ with NH_4_Cl, MgCl_2_, and FeCl_3_, respectively.

### 2. Isolation of gene-disrupted mutants and transconjugant

Targeted gene disruption of *R. capsulatus* was performed using the homologous recombination technique, and all gene cloning operations were performed using an In-Fusion® HD Cloning Kit (Takara Bio, Shiga, Japan). A conjugated pZJD29a vector [65] was employed to clone 500-bp upstream and downstream regions of the target gene. The *islAB* genes, which encode subunits of anaerobic isethionate lyase from *Bilophila wadsworthia* 3_1_6, were synthesized with codon optimization for *E. coli* (Invitrogen GeneArt, Thermo Fischer Scientific, Waltham, MA) and the nucleotide sequence of *islAB* used in this study can be downloaded from https://github.com/Yoshiki-Mo/supporting_data_2024.5. For *islA* and *islB,* an 11-bp intergenic region between *trpB* and *trpC*, present in the tryptophan synthesis operon of *E. coli,* was inserted into the intergenic region [66]. The expression of the *isrBHDK* genes (rcc02237-rcc02236-rcc02235-rcc02234) and the *islAB* genes were regulated by the *pucB* promoter that was inserted just upstream of the initial codons of *isrB* and *islA*. The chimeric gene fragments were then cloned into pBBR-MSC2 [67]. For the expression of *ssuCAB* (rcc02223-rcc02224-rcc02225) or *isrBHDK* in *R. sphaeroides*, the 280-bp upstream sequence of *pucB* from *R. sphaeroides* was employed as a promoter, and the *isrBHDK* or *ssuCAB* genes were inserted downstream of the promoter unit and then cloned into pBBR-MSC2. In addition, plasmids for co-expression of *isrBHDK* and *ssuCAB* genes were constructed by tandem cloning of the respective expression gene regions just downstream of the *pucB* promoter into pBBR-MSC2. Each plasmid was transformed into cells of *R. capsulatus* and *R. sphaeroides* using the conjugative transfer method with *E. coli* S17-1 *λpir*, as described in previous studies [9]. The primer sequences used in this work are listed in Table S4.

### 3. Analysis of genes specifically induced by isethionate under photosynthetic conditions

#### 3-1. RNA extraction and RNA-seq library preparation

*R. capsulatus* cells were grown in RCV-SUL or RCV-ISE medium under anaerobic photosynthetic conditions. Total RNA was extracted from cells in the exponential phase (OD_660_ = 0.2-0.3; UV-1800, Shimadzu, Kyoto, Japan). For RNA preparation, 1 mL of cells was mixed with 200 µL of Stop solution (5% phenol: 95% ethanol), vortexed for a few seconds, and then centrifuged quickly (13,300 rpm, AR015-24, TOMY, Tokyo, Japan). Cells were resuspended in TE buffer (100 µL) containing 10 mg/mL lysozyme and incubated at room temperature for 5 min. RNA was extracted using NucleoSpin RNA (Macherey-Nagel, Düren, Germany) according to the manufacturer’s protocol. Genomic DNA was on-column digested with RNase-free DNase treatment. The concentration of total RNA was determined by measuring the absorbance at 260/280nm using a NanoDrop Spectrophotometer (Thermo Fischer Scientific). Library preparation and stranded RNA-seq were performed by Genome-Lead (Takamatsu, Japan). In brief, the MGIEasy RNA Directional Library Prep set (MGI) was used for reverse transcription and subsequent cDNA library preparation. Sequencing was performed using the DNBSEQ-G400RS sequencer, and the 26-41 million sequencing reads obtained from each replication were used for subsequent RNA seq data analysis (Table S5).

#### 3-2. RNA seq

Demultiplexed raw sequencing reads were quality trimmed and adapter filtered using fastp [68] with default settings. Quality filtered reads were aligned to the *R. capsulatus* SB1003 genome (GenBank accession: GCA_000021865.1) using Bowtie2 [69] with global alignment and sensitive settings. SAM alignments were converted to binary sam (BAM) format using the SAMtools sort command [70]. The number of sequencing reads aligned to each open reading frame was counted using FeatureCounts with the option “-p -T 8 -t CDS -g gene_id” (Table S6) [71]. Differential expression gene (DEG) analysis between RCV-SUL and RCV-ISE conditions was performed using the Bioconductor DESeq2 v.1.3.4 package (Figs. S10, S11, Tables S7, S8) [72]. The p-values were adjusted for multiple testing using the Benjamini– Hochberg method to control the false discovery rate (FDR) below 0.001. Genes significantly induced in RCV-ISE conditions with FDR < 0.001 and fold change greater than 100-fold compared with RCV-SUL conditions are shown in Table 1 and Fig. S5. Volcano plots were generated using the ggVolcanoR shiny server (Fig. S11) [73].

### 4. Phylogenetic analysis of NFLs and RSEs

#### 4-1. Search for a single ortholog cluster of Group IV

The determination of orthologs of the various proteins in Group IV NFL was performed using the following two-step procedure (4-1-1 and 4-1-2). Since the Group IV NFLs Isr, Mar, and Nfa comprise the BHDK homolog set and Cfb comprises the HD homolog set, respectively, strains carrying all of the genes in these sets were defined as strains harboring the respective NFLs (Table S9).

##### 4-1-1. Search for homologs

Regarding Isr, homologs of IsrB (WP_013067946.1), IsrD (WP_013067944.1), and IsrK (WP_013067943.1) were BLASTp searched against the NCBI RefSeq Select proteins database with an e-value cut-off of 1e^-10^ and a minimum alignment fraction of 70%. Regarding IsrH (WP_013067945.1), IsrH was retrieved from strains carrying IsrB, IsrD, and IsrK orthologs, as defined in 4-1-2. Homologs of other Group IV NFLs (Cfb, Nfa, and Mar) subunits were searched in two steps. First, the IsrD orthologs were defined as described in 4-1-2. Then, homologs of other subunits (B, H, and K in Mar, Nfa, and H in Cfb) were searched, restricted to species carrying the IsrD ortholog.

##### 4-1-2. Phylogenetic inference of the diverse Group IV NFLs and related sequences

To estimate the phylogenetic relationship of Group IV NFL subunits and evolutionarily related protein families, phylogenetic analysis was performed as follows. First, each homolog sequence set from 4-1-1 was multiple sequence aligned using Mafft version 7.0 with default settings [74]. Multiple sequence alignments were visually checked, and highly gapped positions were trimmed using Clipkit version 1.4.1 with default settings [75]. Maximum likelihood (ML) phylogenetic inference was performed with 1,000 ultra-fast bootstrap replicates and Modelfinder was used to estimate the best-fit substitution model [76] using IQ-TREE version 2.2.0.3 [77]. The resulting tree was visualized using iTOL v.6.0 (Fig. S12) [78]. The monophyletic clade containing known Group IV NFL and related sequences was manually recovered and treated as a single ortholog cluster.

#### 4-2. Phylogenetic inference of the NFL

Phylogenetic analysis of each single ortholog cluster (subunits of NFLs) was iteratively performed on the following sequences; 1) Groups I, II, III, and V NFL orthologs as described in Raymond *et al.,* 2004 [4] and Boyd *et al.*, 2011 [79] and 2) Group IV NFL orthologs explored in 4-1 (Table S9). ModA, which was used as an outgroup in the estimation of the B subunit phylogeny, was obtained from the NCBI UniProtKB/Swiss-Prot database (Table. S10). For ML phylogenetic inference, IQ-TREE was used as explained in the previous section. The treeline used to create the phylogenetic tree can be downloaded from https://github.com/Yoshiki-Mo/supporting_data_2024.5.

### 5. Distribution of isethionate-metabolizing enzymes

#### 5-1. Search for a single ortholog cluster of isethionate-metabolizing enzymes

The homologs of the isethionate-metabolizing enzymes Xsc and IslA were BLASTp searched against the NCBI RefSeq Select proteins database with an e-value cut-off of 1e^-10^ and a minimum alignment fraction of 70%. The top 1,500 hits with zero or the lowest e-value were retrieved. Here, according to 4-1-2, sequences from the query-containing monophyletic clade were considered as single ortholog clusters and manually recovered (Table S11).

#### 5-2. Phylogenetic Inference of strains carrying isethionate metabolizing enzymes

The phylogenetic relationship among strains containing isethionate metabolizing enzymes was inferred using SSU 16S rRNA sequences. Full-length 16S rRNA sequences were obtained using Barnap version 0.9 (https://github.com/tseemann/barrnap) from each genome assembly or manually extracted from the LPSN website (October 17 in 2023 access) [80]. A 1,000 bootstrap replication ML tree estimation was performed according to method 4-1-2. The treeline used to plot the phylogenetic tree is available at https://github.com/Yoshiki-Mo/supporting_data_2024.5.

### 6. Comparative genome analysis for genome profiling

The genome sequences of 14 *R. capsulatus* and 29 *R. sphaeroides* strains were downloaded from the NCBI assembly (December 13, 2022 access). Genomes satisfying Completeness > 90% and Contamination < 5% for 120 conserved proteins according to the MIMAG (meta)genomic criteria were selected [81]. The remaining 41 high-quality (HQ) genomes were confirmed for genome taxonomy using two methods. First, the ANI distances of the 41 genomes against *R. capsulatus* SB1003 (GCA_000021865.1) and *R. sphaeroides* type strains (GCA_000012905.2) were checked using Pyani version 0.2.11 [82]. Second, the phylogenetic placement of 41 HQ genomes was investigated using the genome taxonomy tool kit GTDB-tk version 2.3.0 [83, 84]. Using these two methods, eight strains were classified as *R. capsulatus*_B, five as *R. capsulatus*, and 17 as *Cereibacter_*A *sphaeroides*. The remaining 30 genome sequences (Fig. S7, Tables S1, S2) were annotated using prokka with default settings (https://github.com/tseemann/prokka). The resulting GFF files were used to search for core genes specifically conserved in all of the eight *R. capsulatus_*B strains that retained the Isr genes but not conserved in the 17 *C_*A*. sphaeroides* strains and the five *R. capsulatus* strains that did not retain the Isr gene.

Pan-genome analysis was performed using PIRATES with “-s 60, 70, 80, 90, 95 -a” option [85]. Scoary package was employed to identify shared genes within a species of *R. capsulatus*_B containing Isr, which is assumed to possess metabolic capability for isethionate [86]. Statistical significance was determined at p < 0.01 after applying Bonferroni multiple test correction (Table S12).

## Acknowledgements

We thank Jiro Harada in Kurume University for providing *R. sphaeroides* J001 and technical suggestions for cultivation. We thank Shinji Masuda in Tokyo Institute for Technology for providing pZJD29a plasmid. We thank Asako Segawa for technical assistance, and Mari Banba and Takafumi Yamashino and all members of the Laboratory of Molecular and Functional Genomics in Nagoya University for engaging in discussion and offering technical assistance.

## Author Contributions

**Haruki Yamamoto:** Conceptualization; data curation; format analysis, investigation; supervision; resources; project administration; visualization; writing – original draft preparation; writing – review and editing; funding acquisition. **Yoshiki Morimoto:** Data curation; format analysis; investigation; visualization; writing - original draft preparation; writing – review and editing; funding acquisition. **Kazuma Uesaka:** Data curation; format analysis; investigation; software; visualization; writing – review and editing. **Yuichi Fujita:** Conceptualization; supervision; resources; project administration; visualization; writing – original draft preparation; writing – review and editing; funding acquisition.

## Funding

This work was supported by Grants-in-Aid for Scientific Research No. 20K06542 and 24H02075 from the Japan Society for the Promotion of Science (JSPS) to H.Y. and Y.F., respectively, TOYOAKI SCHOLARSHIP FOUNDATION to H.Y., and Nagoya University Interdisciplinary Frontier Fellowship to Y.M.

## Data Availability Statement

Raw RNA-seq sequencing reads have been deposited in the DDBJ Sequence Read Archive (DRA) under BioSample accession numbers SAMD00770669. Data are contained withing the article or Supplementary Material. All raw data have been deposited into the data managing system in Nagoya University.

## Conflicts of Interest

The authors declare no conflicts of interest.

## Notes

https://github.com/Yoshiki-Mo/supporting_data_2024.5

### Competing Interest Statement

The authors have declared no competing interest.

