## supplemental figures for "A nitrogenase-like enzyme is involved in the novel anaerobic assimilation pathway of a sulfonate, isethionate, in the photosynthetic bacterium *Rhodobacter capsulatus*"

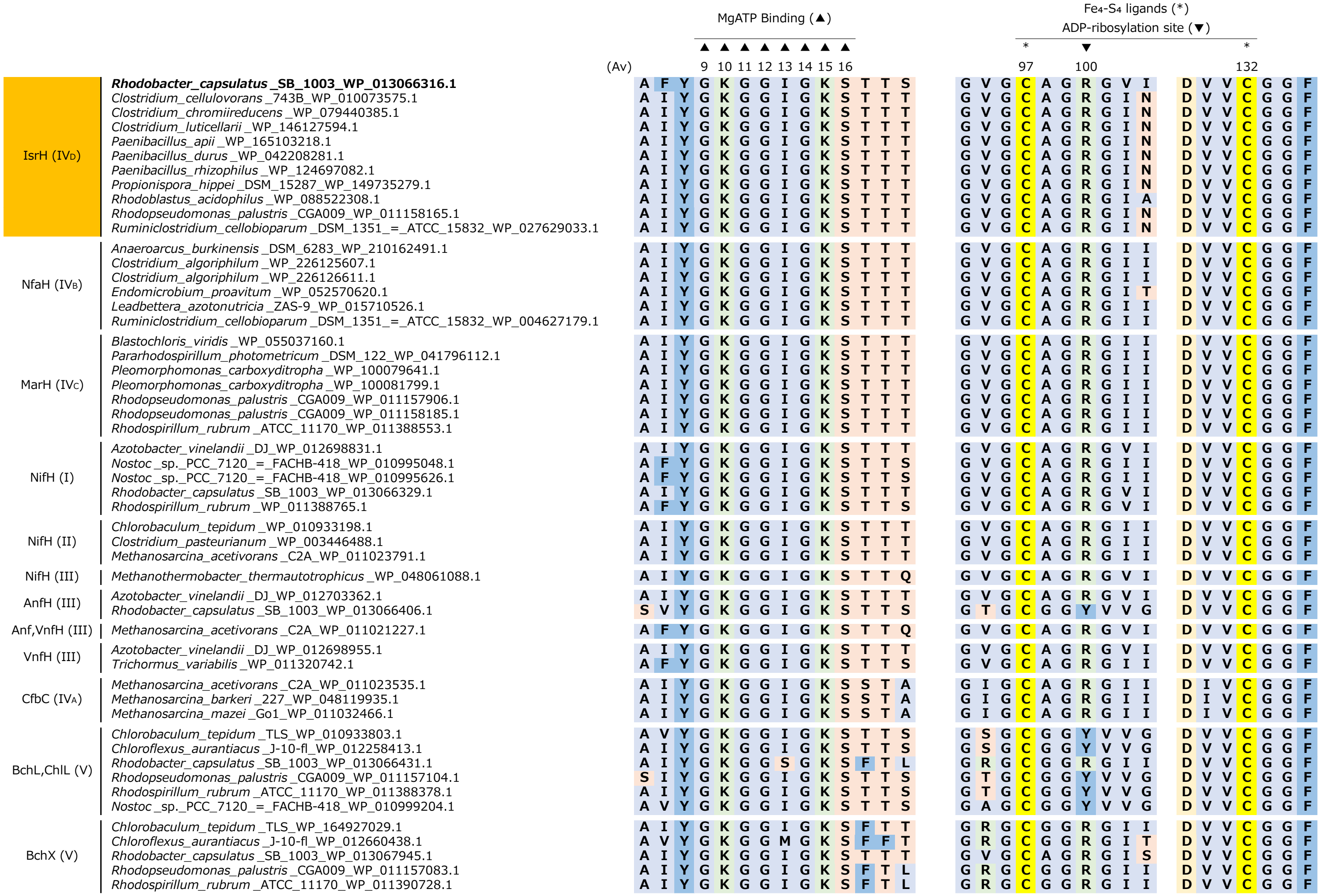


**Supplementary Fig. S1. Partial multiple sequence alignment of the H homologs; IsrH, NfaH, MarH, NifH, AnfH, VnfH, CfbC, BchL/ChlL, and BchX.**

Each ortholog sequence set were multiple sequence aligned with Mafft version 7.0 with default settings. Three motifs for ATP-binding (14 amino acid residues) and the iron-sulfur cluster binding (10 and 7 amino acid residues containing two Cys residues for chelating the [4Fe-4S] cluster) are shown. In the first iron-sulfur cluster motif Arg100 for ADP-ribosylation site in purple bacteria is also shown. (Av) denotes the amino acid sequence number in NifH from *A. vinelandii* DJ (WP_012698831.1.1). Cysteine residues are in yellow (C), acidic amino acids are in light yellow (D, E), basic amino acids are green (H, K, R), neutral polar amino acids are orange (N, S, T, Q), and aromatic amino acids are dark blue (F, W, Y), Amino acid residues with non-polar side chains are shown as light blue (A, G, I, M, V, P, L).

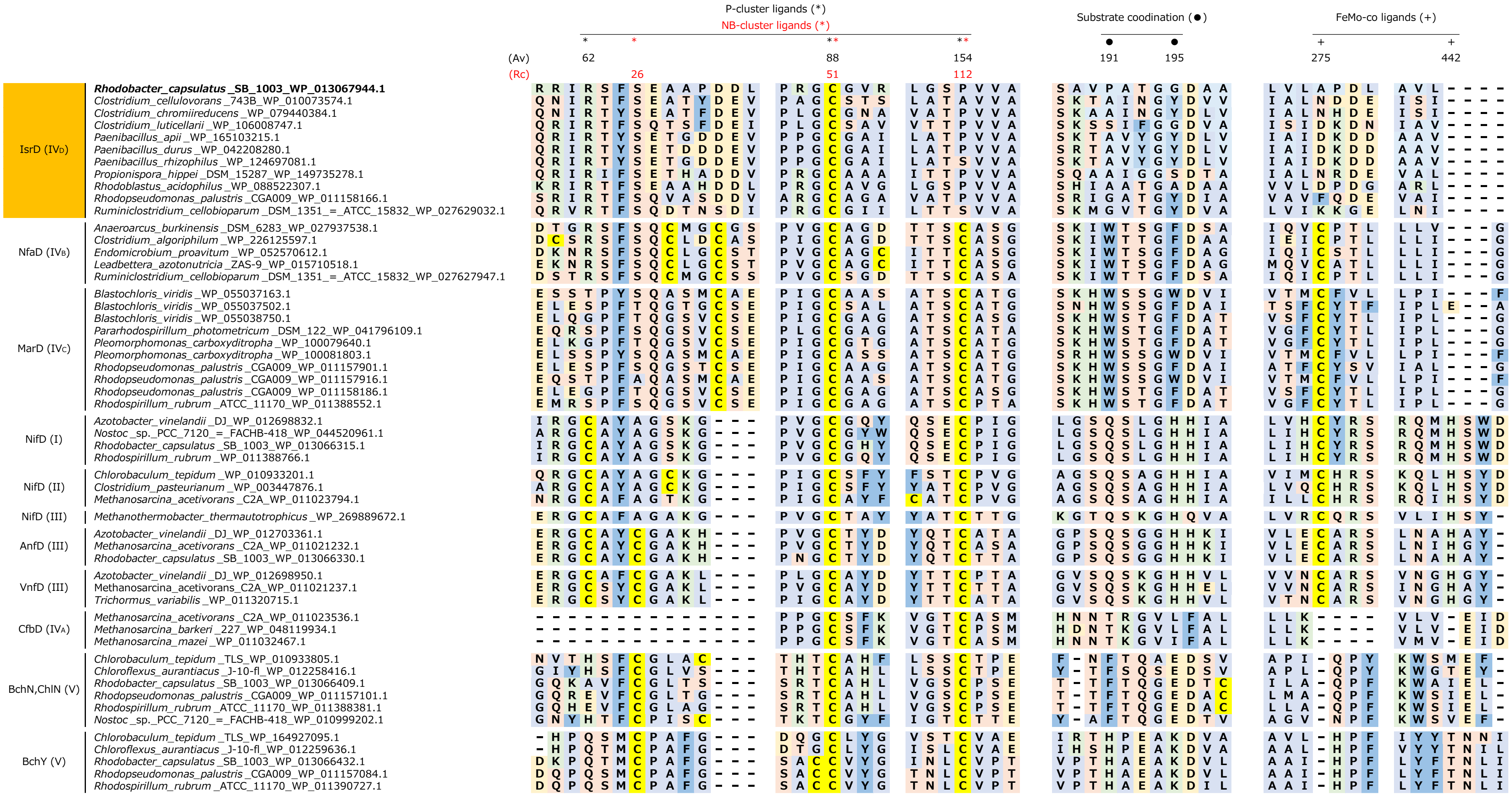


**Supplementary Fig. S2. Partial multiple sequence alignment of the D homologs; IsrD, NfaD, MarD, NifD, AnfD, VnfD, CfbD, BchN/ChlN, and BchY.**

Each ortholog sequence set was multiple sequence aligned in the same manner as Fig. S1. Six motifs for the P-cluster binding (NifD), the substrate binding (BchN), and FeMo-co binding (NifD) are shown. (Av) and (Rc) denote the amino acid sequence numbers in NifD from *A. vinelandii* DJ (WP_012698831.1.1) and BchN from *R. capsulatus* (WP_013066315.1). The color representation of amino acid residues is the same as in Fig. S1.

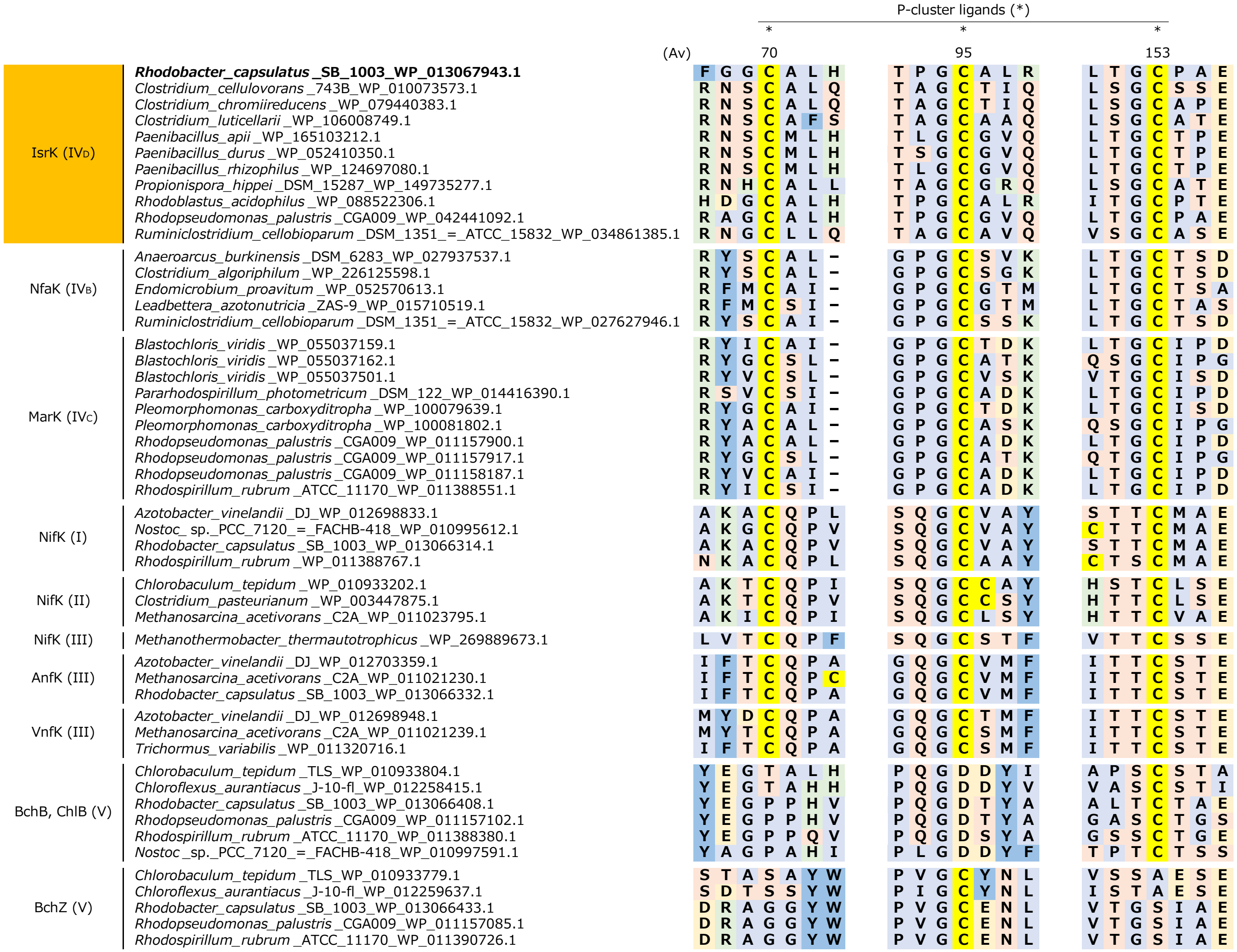


**Supplementary Fig. S3. Partial multiple sequence alignment of the K homologs; IsrK, NfaK, MarK, NifK, AnfK, VnfK, BchB/ChlB, and BchZ.**

Each ortholog sequence set was multiple sequence aligned in the same manner as Fig. S1. Three motifs containing three Cys residues for the P-cluster chelating in NifK are shown. (Av) denotes the amino acid sequence numbers in NifK from *A. vinelandii* DJ (WP_012698833.1.1). The color representation of amino acid residues is the same as in Fig. S1.

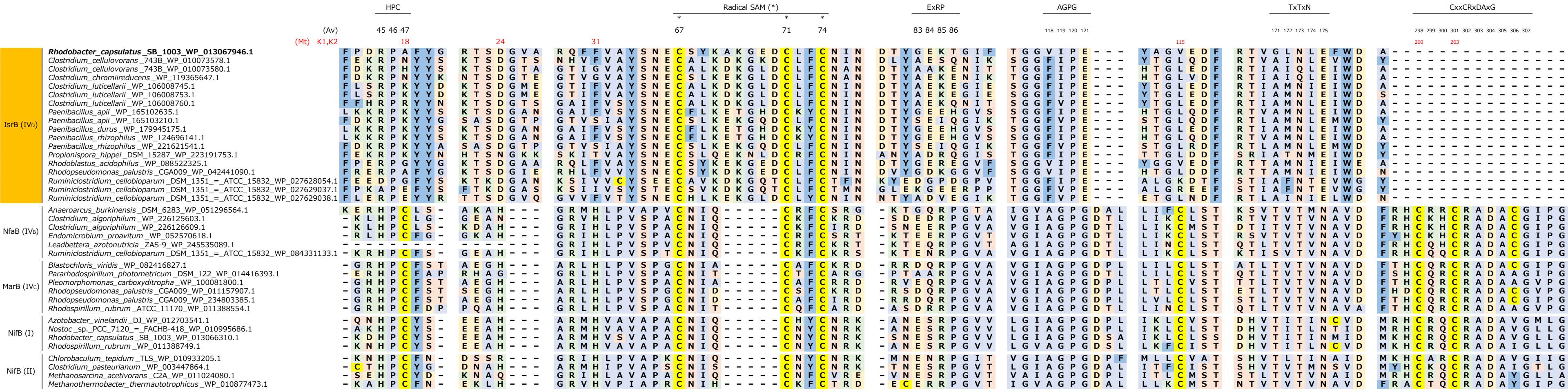


**Supplementary Fig. S4. Partial multiple sequence alignment of the B homologs; IsrB, NfaB, MarB and NifB.** Each ortholog sequence set were multiple sequence aligned with Mafft version 7.0 with default settings. Seven motifs including the radical SAM motif are shown. (Av) and (Mt) denote the amino acid sequence numbers in NifBs from *A. vinelandii* DJ (WP_012703541.1) and *Methanothermobacter thermautotrophicus* (WP_010877473.1), respectively. The color representation of amino acid residues is the same as in Fig. S1.

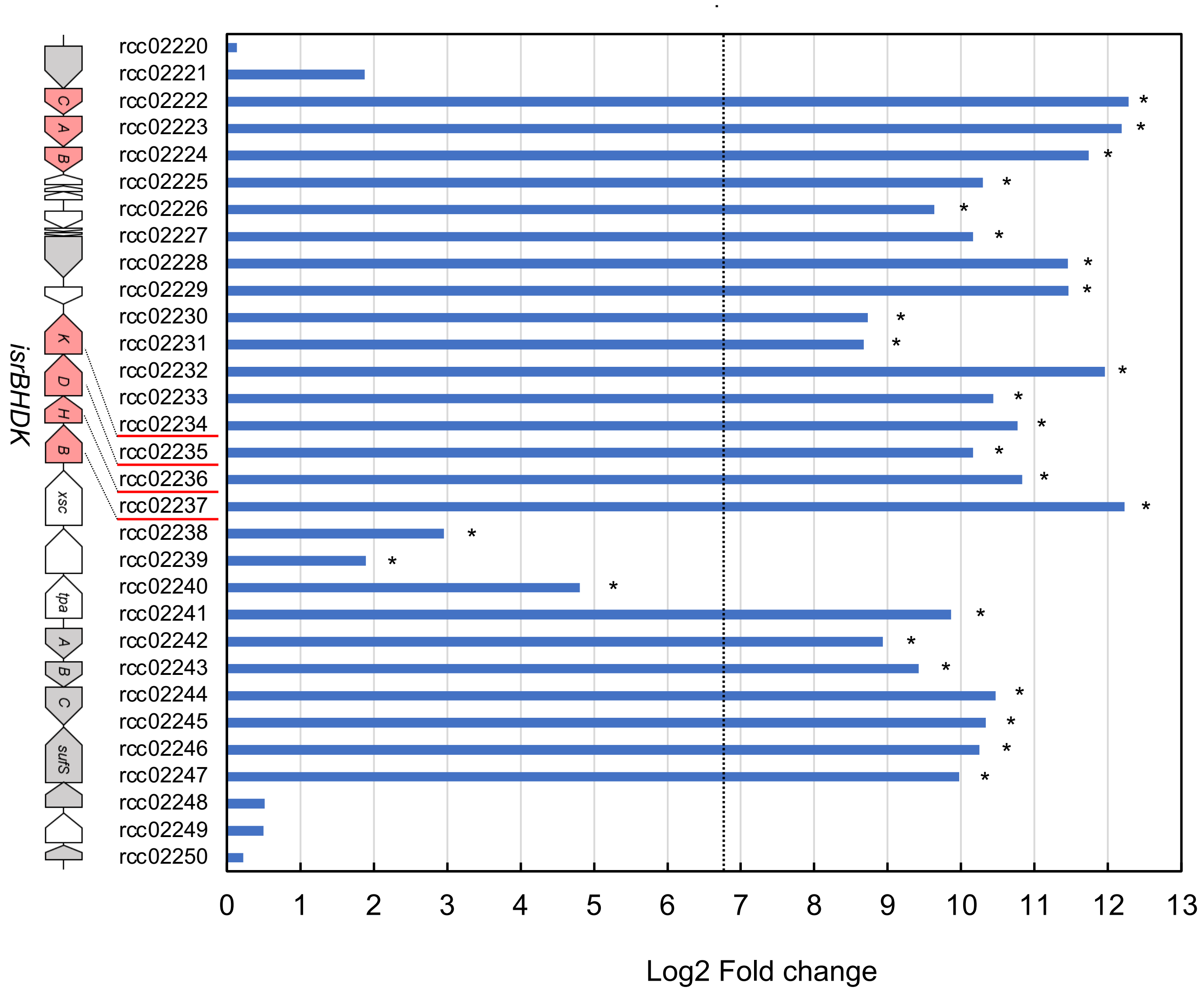


**Supplementary Fig. S5. Genes significantly induced in medium containing isethionate as the sole sulfur source**

The Ise/Sul ratios (the transcript levels in the medium containing isethionate to that in the medium containing sulfate as the sole sulfur sources) for the 31 genes (rcc02220–rcc02250) in the 26-kb gene cluster are shown as values of log2 fold change. $Log_2 100$ (=6.64) (indicated by the dashed line) is the criterion for an Ise/Sul ratio greater than 100, which is significantly induced in the Ise condition. Genes with FDR<0.001 in the test by DEseq2 were marked with * (Table S8).

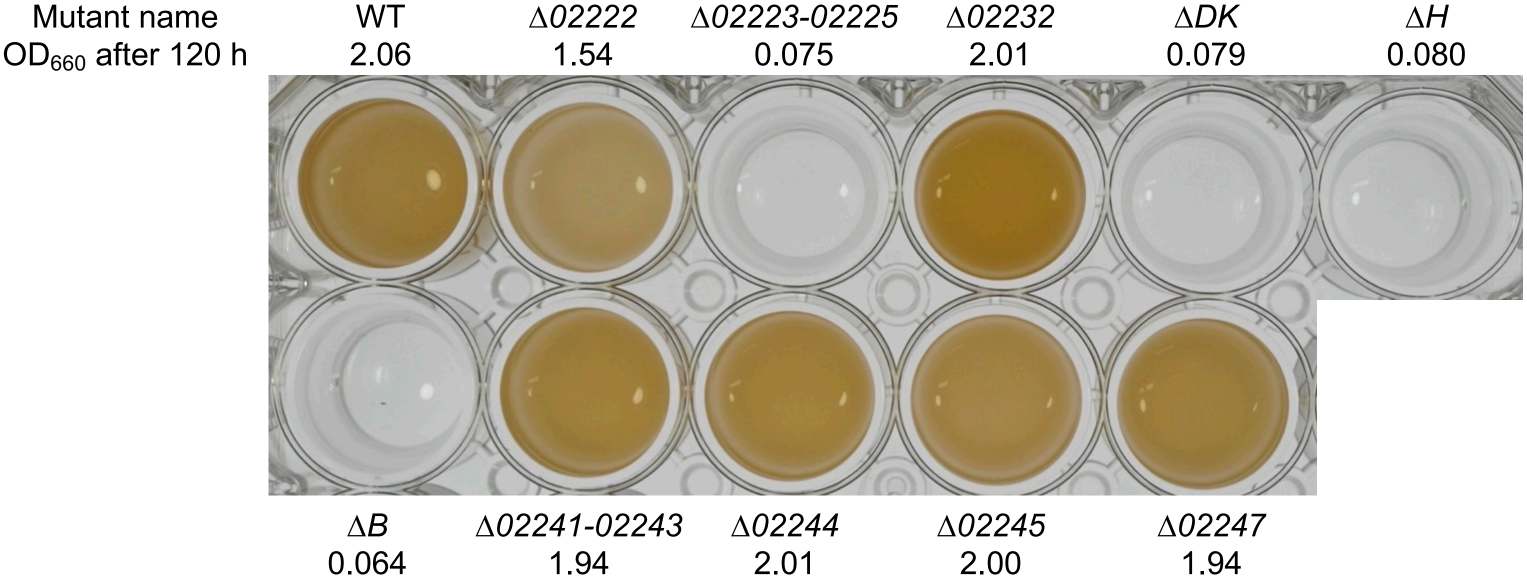


**Supplementary Fig. S6. Growth of ten knockout mutants including three *isr* mutants (Δ*B*, Δ*H*, and Δ*DK*) under anaerobic photosynthetic conditions with isethionate as the sole sulfur source.**
WT and the mutants were grown in test-tubes and transferred to a micro-titer plate as described in Figure 3. OD$_{660}$ values after 120-h incubation are shown below the mutant names.

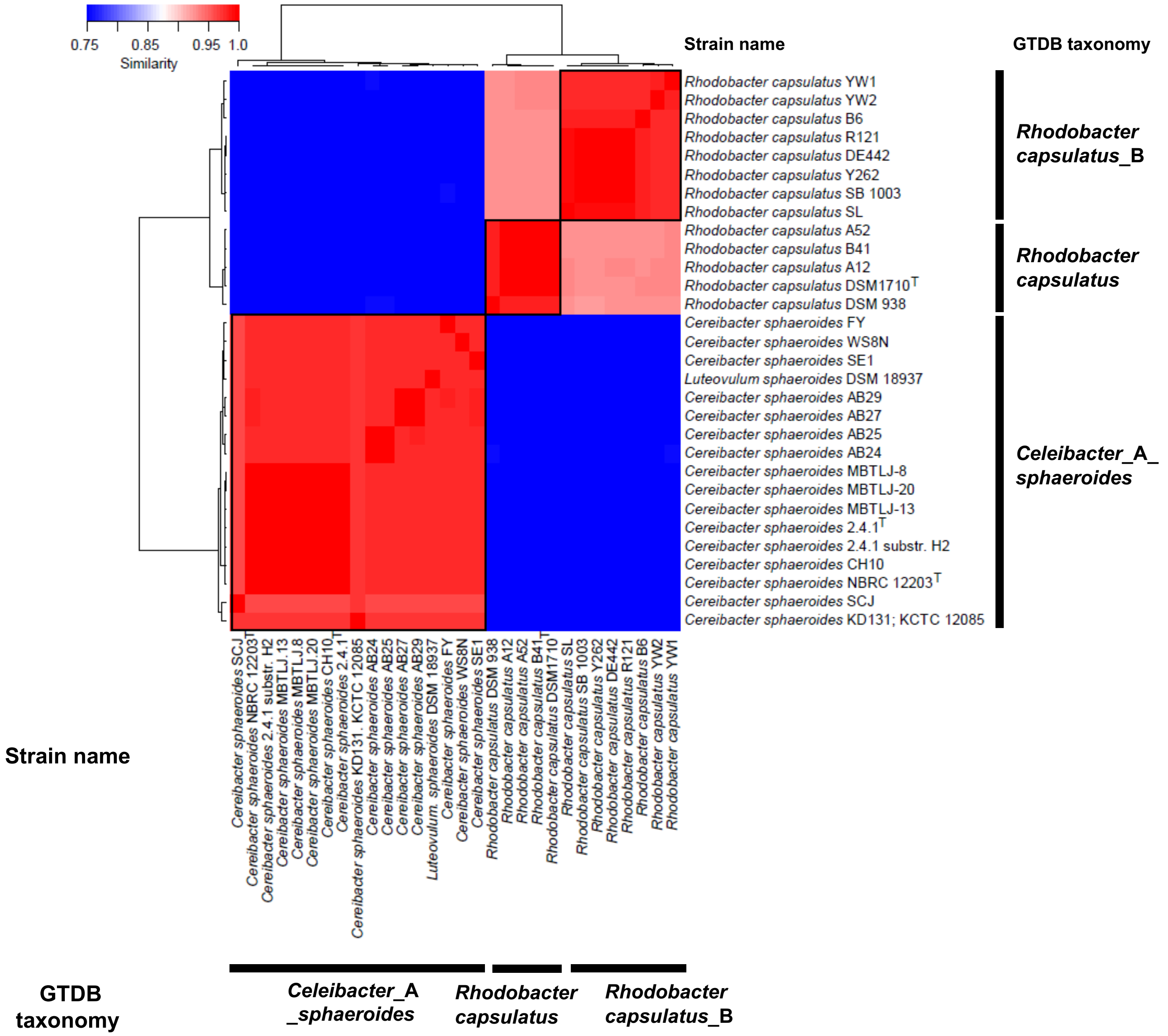


**Supplementary Fig. S7. Evaluation of ANI in 30 strains used for core gene comparisons.**
Heatmap clustered by hierarchical clustering, where T represents the type strain. Clusters with 95% ANI are surrounded by thick black lines.

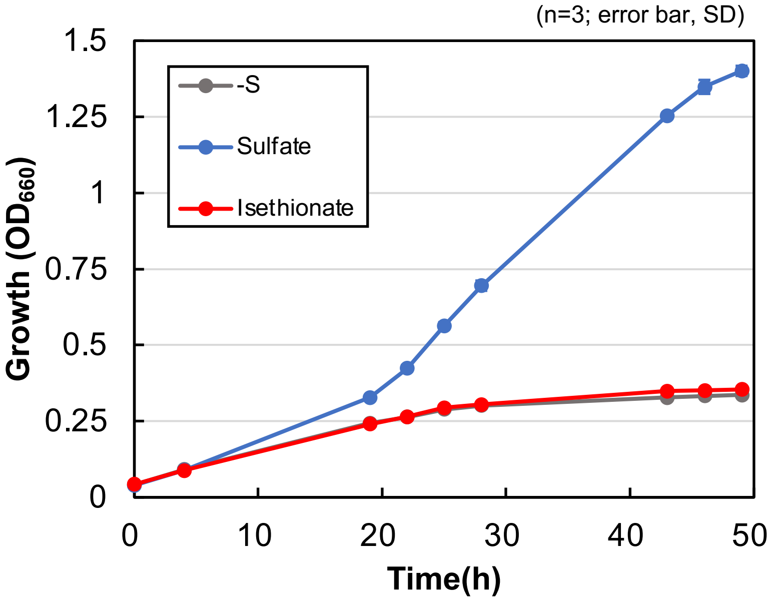


**Supplementary Fig. S8. Growth of *R. sphaeroides* 2.4.1 using sulfate and isethionate as the sole sulfur sources.**

To examine whether *R. sphaeroides* 2.4.1. J001-1 has growth ability using isethionate as the sole sulfur source, *R. sphaeroides* J001-1 was grown in Sistrom medium. Sistrom (-S) was used a negative control for no sulfur medium, and $MgSO_4$ (Sulfate) and sodium isethionate (Isethionate) were added to the Sistrom medium at the final concentration of 0.1 mM, respectively. *R. sphaeroides* J001-1 was grown under anaerobic photosynthetic heterotrophic conditions. Growth was monitored with $OD_{660}$.

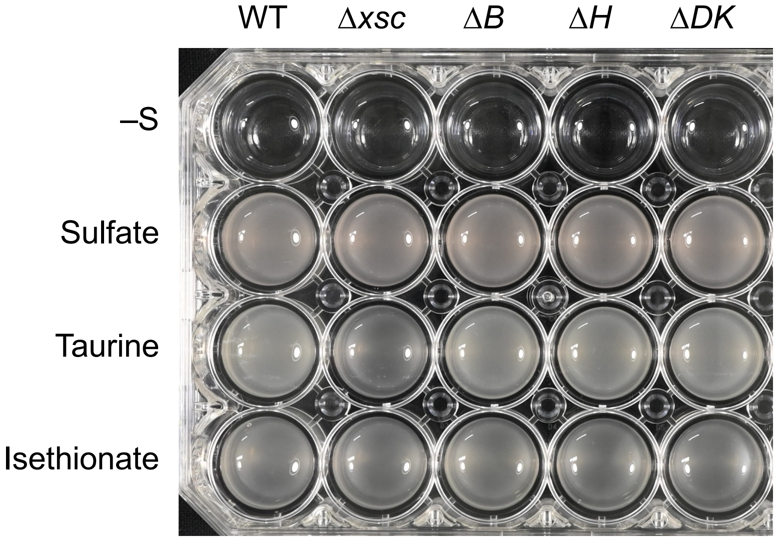


**Supplementary Fig. S9. Growth of the four *isr* mutants of *R. capsulatus* under aerobic heterotrophic conditions with three sulfur sources.**
WT and the mutants were grown with sulfate, taurine, and isethionate as sulfur sources in test tubes under aerobic conditions (reciprocal shaking at 105 rpm; 34ºC, 96 h), and the cultures were placed in a 24-well plate to show clearly their growth. Note that the final turbidity of even WT when grown under aerobic respiration conditions is much lower than that of growth under anaerobic photosynthetic conditions (Fig. 3).

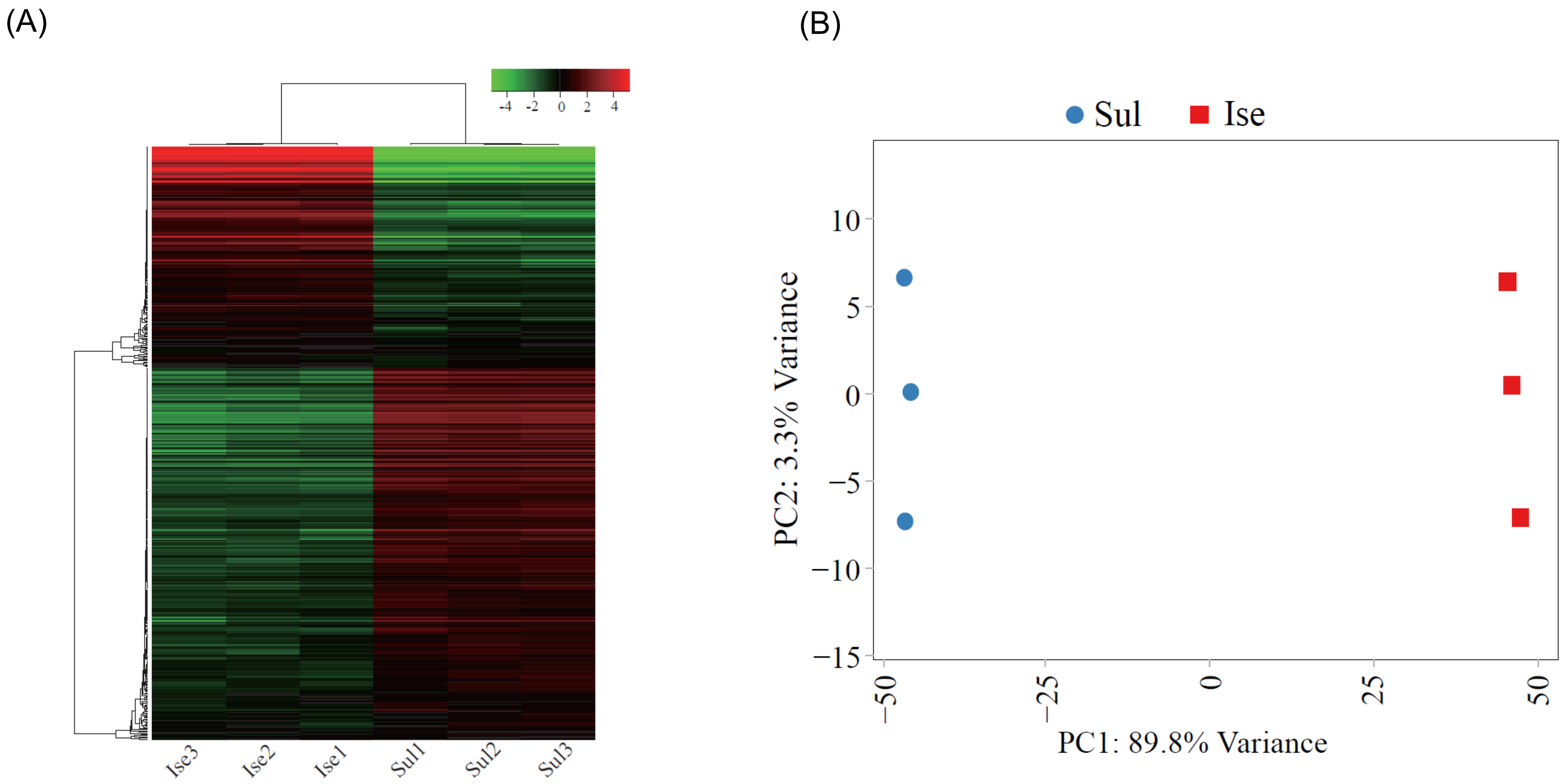


**Supplementary Fig. S10. RNA-seq quality control metrics.**
(A) Unsupervised hierarchical clustering heatmap. Heatmap was generated based on z-scaled Log CPM (Counts Per Million) expression counts between two groups, with 3 replicates each. (B) Principal component analysis (PCA) of RNA-seq data between Ise and Sul shows major separation of samples by the sulfur source.

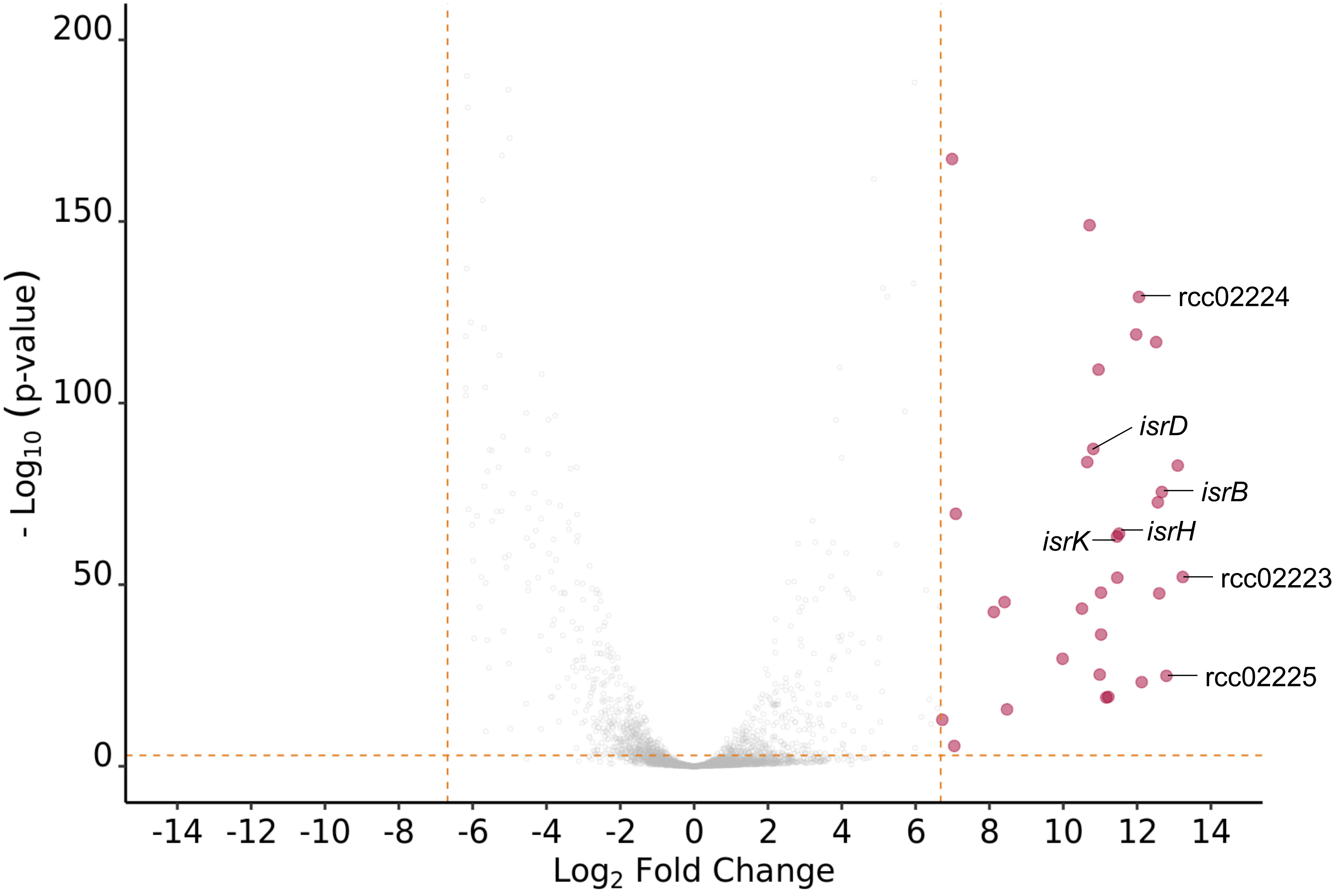


**Supplementary Fig. S11. Volcano plot displaying differentially expressed genes in Ise and Sul cells.**
Red dots indicate genes with significantly different expression levels in the Ise condition. Y-axis represents -$\log_{10}$ FDR (False Discovery Rate), and the x-axis represents $\log_2$ fold change values. Dashed lines indicate cut-off borders. All genes shown in Fig. S5 as significantly induced under Ise conditions are included in this figure. The plots corresponding to the seven genes, *isrBHDK* and *ssuCAB* (rcc02223, rcc02224, and rcc02225) are indicated with the gene names.

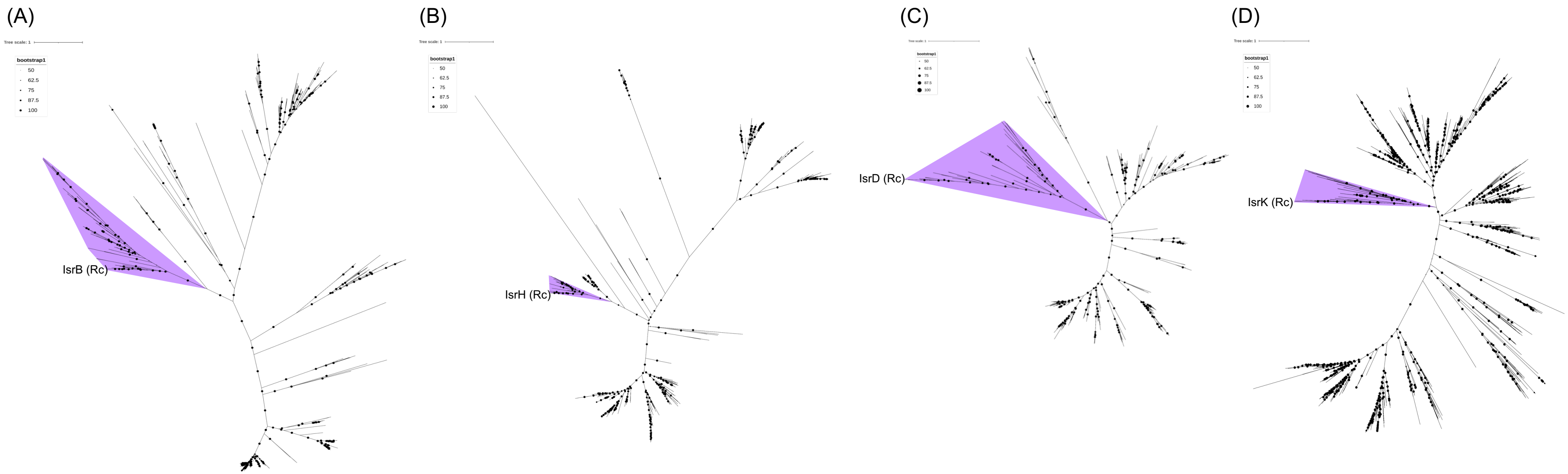


**Supplementary Fig. S12. Molecular phylogenetic trees used to extract the *isrBHDK* orthologs from various bacterial species.**
Homologs to IsrB (A), IsrH (B), IsrD (C), and IsrK (D) were extracted as described in Materials and Methods (4-1-1), and molecular phylogenetic trees were constructed (4-1-2). Homologs in the clade containing the Isr subunit of *R. capsulatus* was selected as Isr orthologs (shaded with purple). Clades were determined based on Bootstrap values and length of branches from the common ancestor. The numbers of IsrB, IsrH, IsrD, and IsrK orthologs extracted were 65, 34, 38, and 40, respectively.
